# Dual regulation of the unfolded protein response by IGF2BP3 during ER stress

**DOI:** 10.1101/2025.03.29.646117

**Authors:** Aleksandra S Anisimova, Harald Hornegger, Irmgard Fischer, Gijs A Versteeg, Stefan L Ameres, G Elif Karagöz

## Abstract

Misfolded protein accumulation in the endoplasmic reticulum (ER) perturbs cellular homeostasis, causing pathological ER stress. The Unfolded Protein Response (UPR) is a highly conserved signaling cascade that restores ER homeostasis by countering ER protein overload. Transcriptional response is paramount for UPR signaling and negating ER stress. While multiple UPR-linked mRNAs are post-transcriptionally regulated, the mechanisms mediating this regulation are unclear. Here, we demonstrate that the highly conserved RNA-binding protein IGF2BP3 interacts with transcripts encoding a subset of UPR effectors. During ER stress, IGF2BP3 destabilized many of these target transcripts, including UPR targets. In contrast, IGF2BP3 stabilized mRNAs encoding transcriptional regulators and thereby upregulated expression of UPR target genes. This dual regulation allows IGF2BP3 to differentially upregulate stress response genes while tuning down the expression of other transcripts during ER stress, relieving protein folding load during this critical response. Our data reveal that posttranscriptional mechanisms control transcription, thus forming gene regulatory networks that robustly tune the UPR.

## Introduction

Protein-folding homeostasis is paramount for robust cell function. Underlining the importance of this homeostasis, various stress-response pathways surveil and react to protein-folding perturbations. In the endoplasmic reticulum (ER), a critical site for the folding of secreted and transmembrane proteins, protein-folding homeostasis is maintained by a conserved signaling cascade known as the “Unfolded Protein Response” (UPR). In metazoans, the UPR is driven by three parallel ER-tethered sensor/transducers: IRE1, PERK, and ATF6. Each sensor monitors the accumulation of misfolded proteins in the ER and takes corrective actions to restore homeostasis.

The UPR increases the capacity for protein folding and degradation in the ER through transcriptional upregulation of chaperones, foldases, ER-associated degradation (ERAD), and ER-phagy components ^1-11^. However, if homeostasis is not achieved in a timely manner, the UPR initiates cell death programs that eliminate defective cells for the benefit of the organism ^12, 13^. Together with the magnitude and duration of the ER stress, the interplay between the UPR branches sets molecular timers that govern the decision to either repair the damaged cell or eliminate it as a potential threat. These factors heavily depend on cell identity as well as metabolic and cellular state ^12, 13^.

The UPR-driven transcriptional response is essential for maintaining ER homeostasis. However, it has become increasingly clear that additional post-transcriptional mechanisms play a crucial role in cellular adaptation to ER stress ^14-18^. The early response to ER stress converges on the shutdown of global translation by the ER-tethered kinase PERK, which reduces global protein synthesis ^19-22^. During translational shutdown, some translationally silenced mRNAs are protected from turnover, whereas others are degraded ^23-25^. Under such conditions, the UPR sensor IRE1 cleaves ER-localized mRNAs via regulated IRE1-<u>d</u>ependent <u>d</u>ecay of messenger RNAs (RIDD), proposed to decrease protein folding load in the ER ^26, 27^. While it is clear that cells regulate mRNA translation and stability during ER stress, how this is mechanistically achieved has been poorly understood.

RNA-binding proteins (RBPs) mediate posttranscriptional mechanisms that facilitate adaptation to developmental and metabolic changes or cellular perturbations ^28, 29^. Using immunoprecipitation-coupled to mass spectrometry analyses (IP-MS/MS) of the UPR sensor IRE1, we previously revealed that it is associated with a highly conserved RBP, IGF2BP3, in a stress-dependent manner ^30^. IGF2BPs are a conserved family of RBPs that regulate mRNA fate by controlling target mRNA stability, localization, and translation (reviewed in ^31^. There are three IGF2BP paralogs in mammalian cells. While IGF2BP1 and IGF2BP3 are expressed at high levels during early development, and their expression levels decrease in adult tissues, IGF2BP2 is expressed at stable levels throughout life ^32, 33^. Notably, expression of all IGF2BP paralogs is upregulated in various cancers, and this strongly correlates with poor patient outcomes, enhanced tumor growth, drug resistance, and metastasis ^32, 34, 35^. Recent data suggest that IGF2BP family members function as m6A and m7G readers. All IGF2BPs guard m6A-modified mRNAs from decay, while IGF2BP1 and IGF2BP3 facilitate the degradation of mRNAs harboring m7G modifications ^36, 37^. Several studies show that during proteotoxic stress induced by heat shock or arsenite treatment, IGF2BPs promote mRNA stability by preventing mRNA degradation. Moreover, they facilitate target mRNA translation once stress is relieved ^36, 38, 39^. Given these data, IGF2BPs may mediate post-transcriptional regulation in response to organelle-specific stress. However, whether IGF2BP3 responds to ER stress, a critical site for protein folding, is unclear.

In this study, we assessed the role of IGF2BP3 in regulating UPR-associated transcripts. Our data confirm that the stress-dependent association of IGF2BP3 with IRE1 is conserved across cell lines and reveal that IGF2BP3 binds prominent UPR-effector transcripts. Using genome-wide transcriptome and mRNA stability analyses, we show that IGF2BP3 regulates the UPR at both the post-transcriptional and transcriptional levels, mediating a failsafe system for robust ER-stress regulation. Our data demonstrate that IGF2BP3-driven posttranscriptional pathways form a unique gene regulatory network that elicits potent UPR.

## Results

### IGF2BP3 localizes to the ER membrane through interaction with ER-bound mRNAs

To decipher posttranscriptional mechanisms regulating mRNA fate during ER stress, we focused on the primary posttranscriptional regulator, the ER-tethered RNase IRE1, which binds to and cleaves the mRNAs crucial for life and death decisions during ER stress. In previous work, we mapped the IRE1 interactome by IP-MS/MS in HEK293T and multiple myeloma cells and identified select RBPs that are associated with IRE1 ^30^. We hypothesized that the RBPs interacting with IRE1 during ER stress regulate the stability or translation of UPR-relevant mRNAs associated with IRE1. Among those RBPs, IGF2BP3 stood out as it was also recovered in IRE1 IP-MS/MS under denaturing conditions after RNA-protein cross-linking, which suggested a robust association with IRE1 and RNA-containing complexes (**Supp. Table 1**, data from ^30^).

To test whether the IRE1-IGF2BP3 interaction extended across different cell types, we performed IPs followed by Western Blot analyses in mouse embryonic fibroblasts without stress and at distinct time points after ER stress induction. IGF3BP3’s interaction with IRE1 increased with ER stress up to 11-fold at 8 h, compared to non-induced controls (**Fig. 1A**). Consistent with this result, we also recovered IRE1 using reciprocal IGF2BP3 pull-downs in HEK293T cells in an ER stress-dependent manner (**Supp. Fig. 1A**).

**Figure 1.**
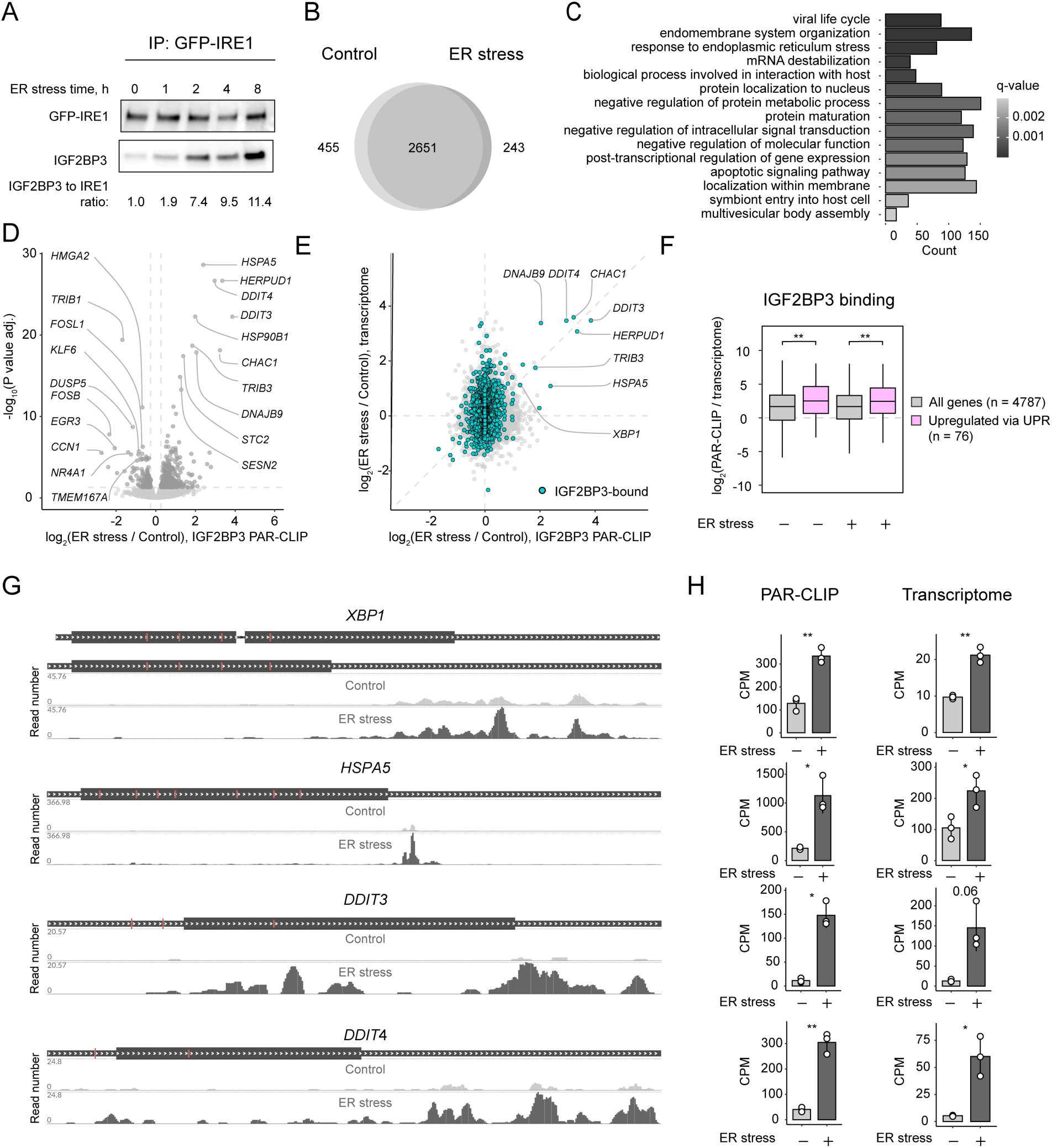
IGF2BP3 interacts with the UPR target mRNAs. **A.** Western blot of IGF2BP3 showing its association with IRE1 after immunoprecipitation of GFP-IRE1 from mouse embryonic fibroblasts treated with ER stress-inducing drug tunicamycin at 5 µg/mL for the indicated time. **B.** Venn diagram showing the intersection of IGF2BP3 IR-PAR-CLIP targets in DMSO control and ER stress conditions (4-hour treatment with 5 µg/mL tunicamycin). **C.** GO term analysis of 2470 IGF2BP3-bound transcripts identified in either control or ER stress conditions with IR-PAR-CLIP CPM higher than transcriptome (QuantSeq) CPM. **D.** Volcano plot showing changes in IGF2BP3 IR-PAR-CLIP CPM upon ER stress treatment. *P* values were calculated by edgeR glmQLFTest. **E.** Scatter plot comparing changes in IGF2BP3 IR-PAR-CLIP read counts with the total transcriptome changes (QuantSeq) upon ER stress treatment. **F.** Boxplot showing changes in IGF2BP3 binding (IR-PAR-CLIP CPM / QuantSeq CPM) to all genes and UPR upregulated genes (2x more in ER stress than in control) upon ER stress and control conditions. *P* values were calculated by two-sided Wilcoxon test. **G.** Representative IGF2BP3 coverage examples and **H.** barplots showing normalized read count numbers (CPM) for IGF2BP3-bound reads (IR-PAR-CLIP) and total transcript levels (QuantSeq) for selected UPR target genes (*HSPA5*, *XBP1*, *DDIT4*, and *DDIT3*). Values are the mean ± s.d. For IR-PAR-CLIP experiment n=3 biological replicates. *P* values were calculated by two-sided Student’s t-test. Where not indicated otherwise, ER stress was induced with tunicamycin at 5 µg/mL for 4 hours. \**P* < 0.05; \*\**P* < 0.01; \*\*\**P* < 0.001; \*\*\*\**P* < 0.0001.

From these results, we concluded that IRE1 and IGF2BP3 form complexes in cells, suggesting that IGF2BP3 interacts with IRE1 mRNA targets and is partially localized to the ER membrane during stress. The fact that these complexes were identified in multiple cell types makes it likely that they are universal cellular complexes with potential importance for general cellular biology.

### IGF2BP3 interacts with the UPR target mRNAs

The mRNAs interacting with IGF2BP3 under steady-state conditions have been well established ^40-42^, but it is unclear whether proteotoxic stress modulates IGF2BP3 interactions with target mRNAs. To this end, we defined IGF2BP3-interacting mRNAs during ER stress in HCT116 cells using Infra-Red Photoactivatable Ribonucleoside-enhanced Crosslinking and Immunoprecipitation (IR-PAR-CLIP) ^40^. This approach allows nucleotide-resolution detection of RBP-interaction sites in target RNAs ^42^. IGF2BP3 IR-PAR-CLIP was performed under homeostatic conditions, during early response (1 h), and at peak response (4 h) time points after UPR induction by tunicamycin (TM). As UPR target genes are transcriptionally upregulated under these conditions, we performed total transcriptome sequencing in parallel to account for changes in transcript levels.

Under homeostatic conditions, IR-PAR-CLIP identified 3106 IGF3BP3-bound transcripts (**Fig. 1B, Supp. Table 1**). A short ER stress induction (1 hour) did not result in significant changes in the levels of the UPR-target transcripts or IGF2BP3 binding (**Supp. Fig. 1B**). However, upon exposing cells to 4 hours of ER stress, we observed a potent induction of the UPR. Under these conditions, IGF2BP3 bound a similar set of transcripts as the control conditions, with 243 additional transcripts that only interacted during ER stress (**Fig. 1B**). After correction for transcript levels (IR-PAR-CLIP CPM > transcriptome CPM), we identified 2470 IGF2BP3-bound transcripts in either control or ER stress conditions (**Supp. Table 1**). Gene ontology analyses of those mRNAs revealed that the mRNA metabolism, intracellular communication, and response to ER stress were the top enriched biological processes (**Fig. 1C**).

In agreement with the published data, our IGF2BP3 IR-PAR-CLIP analyses showed that under homeostatic conditions, IGF2BP3 binds to most of its target mRNAs through their 3’ UTRs (**Supp. Fig. 1C**) ^40, 42^. While the preference for 3’ UTR binding was not affected by ER stress, the IGF2BP3-binding motifs showed distinct differences (**Supp. Fig. 1D**). The top five 5 nt-long motifs enriched in IGF2BP3-bound IR-PAR-CLIP reads included those with CA-rich sequences common among IGF2BP3 binding motifs ^41, 43^. Three motifs (UCCAG, AGCCU, and UGCCA) were common for both conditions, while upon stress novel ACUGU and ACCUG motifs were highly enriched in comparison to a canonical CAUU-containing motif ^42^. These results indicated that the binding site sequences associated with IGF2BP3 change during ER stress.

We next tested whether ER stress modulates IGF2BP3 interactions with target transcripts. Notably, the number of IGF2BP3-bound reads originating from UPR-induced transcripts increased upon ER stress induction, while IGF2BP3 bound less strongly to some of its canonical targets, such as *HMGA2* mRNA ^44^, under these conditions. These UPR-induced targets included transcripts encoding for the master transcription factors XBP1, ATF4, CHOP (*DDIT3*), and major chaperones BiP (*HSPA5*) and DNAJB9 (**Fig. 1D**). To account for changes in the mRNA levels, we next compared ER stress-induced changes in transcript levels with the changes in IGF2BP3-bound transcripts (**Fig. 1E**) and calculated an “IGF2BP3 binding” score by normalizing the IR-PAR-CLIP reads to the transcript levels (**Fig. 1F, Supp. Fig. 1E**). These analyses showed that IGF2BP3 already binds UPR-induced targets under homeostatic conditions. However, while IGF2BP3-bound reads increased linearly for multiple UPR targets with increased expression, mRNAs such as *HSPA5* showed preferential binding of IGF2BP3 during ER stress (**Fig. 1E, G, H, Supp. Fig. 1E**). Moreover, we identified novel IGF2BP3 targets, such as *DDIT3,* which were only expressed during ER stress (**Fig. 1G, H**).

Supporting our IP-MS/MS data, IGF2BP3 was bound to one-third of the transcripts interacting with IRE1 during ER stress (**Supp. Fig. 1F**). The IGF2BP3-binding sites did not overlap with IRE1 RIDD cleavage sites on most of the common IRE1 and IGF2BP1-binding RNAs. Those analyses indicated that IGF2BP3 binding does not protect those transcripts from RIDD (**Supp. Fig. 1G**). To sum up, we found that IGF2BP3 strongly interacts with transcripts encoding UPR target genes during ER stress, identifying IGF2BP3 as a potential posttranscriptional regulator of the UPR.

### IGF2BP3 depletion dampens UPR signaling

Based on our findings above, we hypothesized that IGF2BP3 might post-transcriptionally regulate UPR-induced mRNAs. We, therefore, set out to identify which transcripts might be functionally regulated by IGF2BP3. To this end, IGF2BP3 was depleted in colon carcinoma HCT116 cells by CRISPR/Cas9-mediated knockout (**Supp. Fig. 2A**), after which transcriptome analyses were performed under homeostatic and ER-stress conditions (**Supp. Table 2**). IGF2BP3 knockout resulted in moderate changes in the total transcriptome (**Fig. 2A**). Specifically, it led to the downregulation of genes involved in cell proliferation, development and response to growth factors, consistent with previous reports ^35, 40, 45, 46^ (**Supp. Fig. 2B**). To address the effect of IGF2BP3 depletion on the cellular response to ER stress, we defined UPR target transcripts as those upregulated by more than two-fold upon ER stress (**Fig. 2B, Supp. Fig. 2C**) and followed their levels upon IGF2BP3 depletion. Transcriptome analysis showed that the levels of the UPR target transcripts were lower in IGF2BP3 knockouts compared to the parental cells during ER stress (**Fig. 2B, C, Supp. Fig. 2C, D**). While IGF2BP3 depletion had only a moderate impact on individual UPR effector transcripts (**Fig. 2B, Supp. Fig. 2C**), changes were evident when analyzed for the whole group (**Fig. 2B, C, Supp. Fig. 2C, D**). In contrast, the levels of IGF2BP3-bound transcripts that are not UPR effectors were affected to much lesser extent upon IGF2BP3 CRISPR/Cas9-mediated knockout. Thus, IGF2BP3 specifically impacts UPR target transcripts during ER stress (**Fig. 2A, C**).

**Figure 2.**
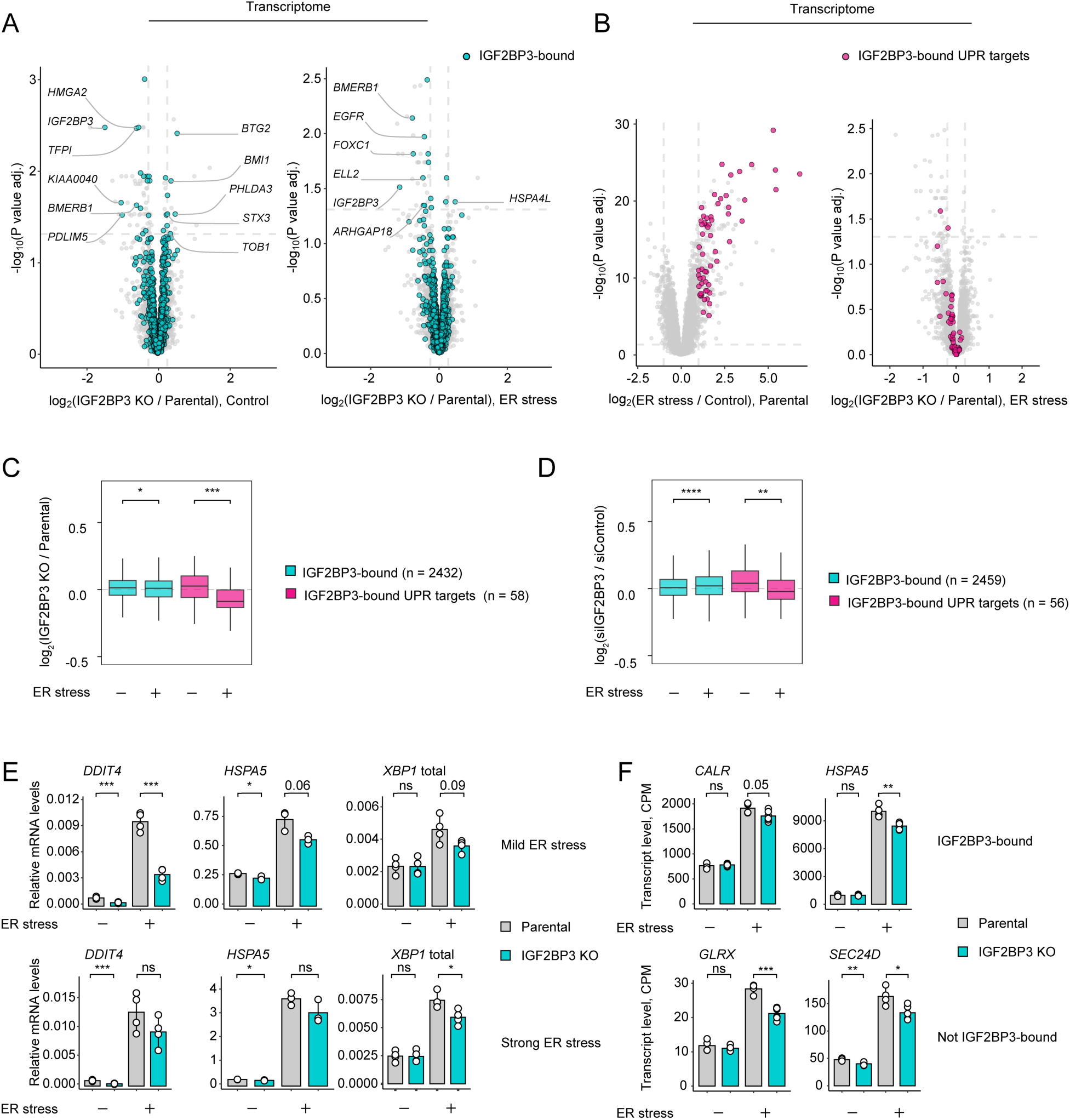
IGF2BP3 depletion dampens the unfolded protein response. **A.** Volcano plot of transcriptome (RNA-seq) changes upon IGF2BP3 KO in control (left panel) and ER stress conditions (right panel). *P* values were calculated by edgeR glmQLFTest. **B.** Volcano plot of transcriptome (RNA-seq) changes upon ER stress treatment showing the selection of IGF2BP3-bound UPR targets (left panel), and volcano plot upon IGF2BP3 KO in ER stress conditions with highlighted IGF2BP3-bound UPR targets (right panel). *P* values were calculated by edgeR glmQLFTest. **C.** Boxplot showing changes in transcript levels upon IGF2BP3 KO for IGF2BP3-bound transcripts and IGF2BP3-bound UPR targets. *P* values were calculated by two-sided Wilcoxon test. **D.** Boxplot showing changes in transcript levels upon siRNA-mediated IGF2BP3 for IGF2BP3-bound transcripts and IGF2BP3-bound UPR targets. *P* values were calculated by two-sided Wilcoxon test. **E.** RT-qPCR analyses of IGF2BP3 KO and parental cells treated with 50 ng/mL (mild ER stress) or 5 µg/mL (strong ER stress) for 7 hours. n=4 biological replicates. *P* values were calculated by two-sided Student’s t-test. **F**. Barplots showing RNA-seq CPM values for selected IGF2BP3-bound and not IGF2BP3-bound transcripts in IGF2BP3 KO and parental cells in control and ER stress conditions. Values are the mean ± s.d. of n=6 biological replicates (duplicates of three different IGF2BP3 KO clonal cell lines), n=4 biological replicates for parental cell line. *P* values were calculated by two-sided Student’s t-test. \**P* < 0.05; \*\**P* < 0.01; \*\*\**P* < 0.001; \*\*\*\**P* < 0.0001.

To account for the adaptation of cells to long-term depletion of IGF2BP3 and to bypass the clonal selection, we used siRNA-based depletion as an orthogonal approach to validate our findings (**Supp. Table 2**). Using an siRNA pool against IGF2BP3, we obtained 75 percent depletion in 72 hours in HCT116 cells (**Supp. Fig. 2E, F**). GO term analyses revealed that IGF2BP3 depletion resulted in lower levels of transcripts involved in adhesion, differentiation, and cytoskeleton organization (**Supp. Fig. 2G**), confirming the function of IGF2BP3 in regulating cell migration and proliferation ^35, 40, 45, 46^. The siRNA knockdown experiments confirmed the data obtained by the CRISPR/Cas9-based IGF2BP3 knockouts, indicating that depletion of IGF2BP3 dampens UPR signaling (**Fig. 2D, Supp. Fig. 2H**). Reverse transcription quantitative PCR (RT-qPCR) analyses of IGF2BP3-interacting transcripts confirmed these results, showing that IGF2BP3 knockout leads to a decrease in the levels of the UPR targets downstream of PERK (*DDIT4*), IRE1 (*XBP1* spliced), and ATF6 (*HSPA5*) (**Fig. 2E**). Notably, IGF2BP3 depletion showed a more substantial effect on UPR signaling when cells were treated with lower tunicamycin concentrations, revealing that IGF2BP3-driven mechanisms might be more pronounced under mild ER stress that is closer to physiological conditions (**Fig. 2E**). Intriguingly, our analyses showed that during ER stress, depletion of IGF2BP3 not only reduced the levels of IGF2BP3-bound UPR targets but also affected those that are not its direct interactors (**Fig. 2F,G, Supp. Fig. 2C, D, H**). This suggested that IGF2BP3 depletion leads to the downregulation of the entire pathway, possibly due to secondary indirect effects. Altogether, our data showed that IGF2BP3 depletion tunes down UPR signaling components.

### IGF2BP3 depletion dampens the translation of UPR effectors

In addition to mRNA stability and localization, IGF2BP3 regulates the translation of a subset of its target mRNAs, the best studied being the *IGF2* mRNA ^47^. To reveal the transcripts whose translation is regulated by IGF2BP3, we performed Ribo-Seq experiments upon siRNA depletion of IGF2BP3 under homeostatic or ER stress conditions (**Supp. Table 2**). We calculated ribosome occupancy by normalizing ribosome-protected fragments in Ribo-Seq experiments to levels of the transcript. IGF2BP3 depletion resulted in very moderate translational regulation which was even less evident upon ER stress induction (**Fig. 3A**). However, IGF2BP3 depletion altered ribosomal occupancy on some transcripts, including those encoding E3 ubiquitin ligase TTC3 and deubiquitinase USP31 (**Fig. 3B, Supp. Fig. 2I**) involved in regulation of AKT ^48^ and NF-κB signaling ^49^, respectively. Notably, IGF2BP3 regulated translation of *DDIT3*, which encodes the stress-regulated transcription factor CHOP (**Fig. 3A, B**). IGF2BP3 depletion resulted in lower CHOP translation, indicating a role of IGF2BP3 in regulating its synthesis. The IR-PAR-CLIP data showed a strong association of IGF2BP3 with the coding region of the *DDIT3* mRNA, distinct from IGF2BP3 interaction with the 3’UTR of most target mRNAs (**Fig. 2F**). Altogether, the Ribo-Seq analyses uncovered a distinct set of mRNAs translationally regulated by IGF2BP3.

**Figure 3.**
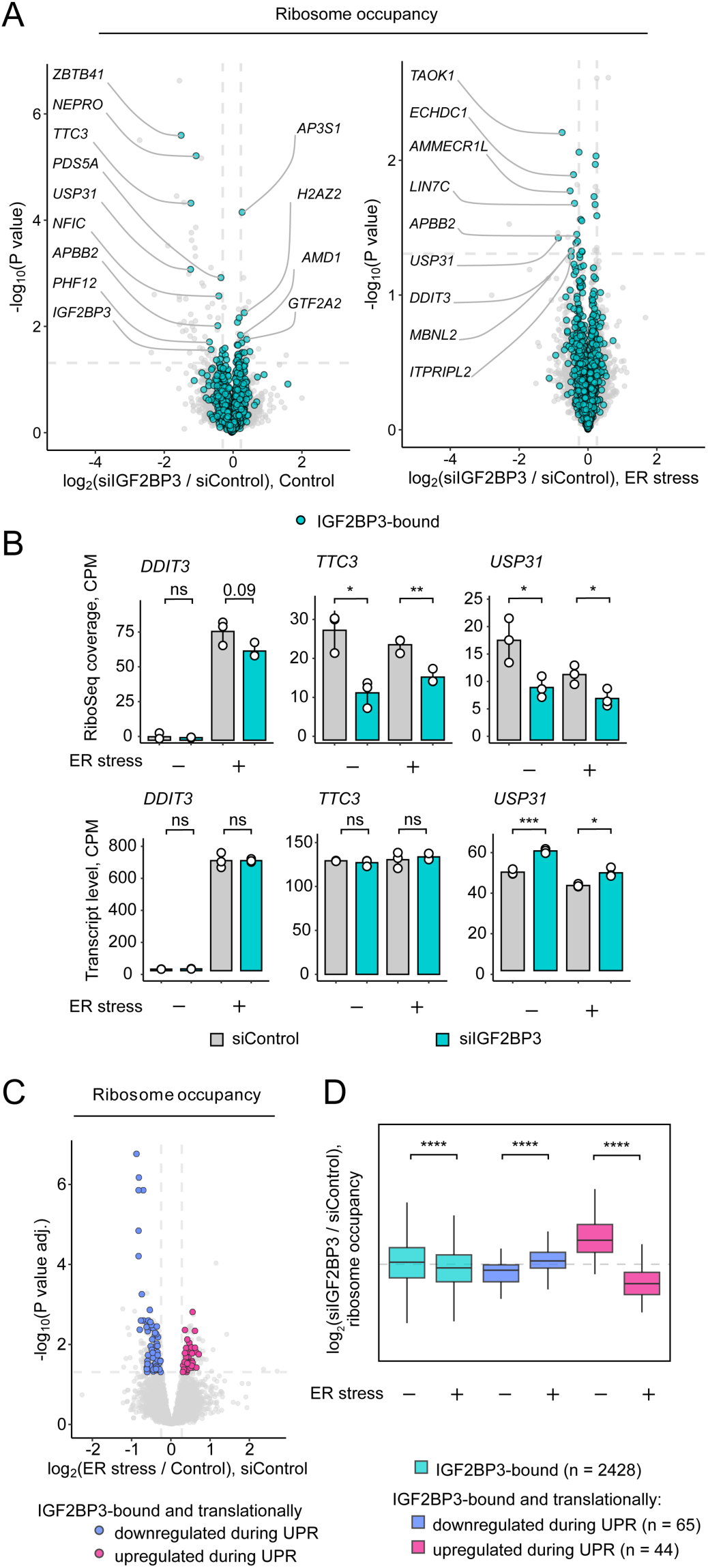
IGF2BP3 depletion dampens the translationally regulated branches of unfolded protein response. **A.** Volcano plots showing changes in ribosome occupancy (RO, log2(RiboSeq CPM / RNA-seq CPM)) upon siRNA-mediated IGF2BP3 depletion in control (left panel) and ER stress conditions (right panel). *P* values were calculated by edgeR glmQLFTest. **B.** Barplots showing RNA-seq and RiboSeq CPM values for *DDIT3*, *USP31*, and *TTC3* in siIGF2BP3- and siControl-treated cells in control and ER stress conditions. Values are the mean ± s.d of n=3 biological replicates. *P* values were calculated by two-sided Student’s t-test. **C.** Volcano plot of ribosome occupancy changes upon ER stress showing the selection of IGF2BP3-bound translationally regulated UPR targets (ΔRO > 20% and *P* value adj. < 0.05). *P* values were calculated by edgeR glmQLFTest. **D.** Boxplot showing changes in ribosome occupancy upon siRNA-mediated IGF2BP3 depletion for IGF2BP3-bound transcripts and IGF2BP3-bound translationally regulated UPR targets. *P* values were calculated by two-sided Wilcoxon test. ER stress was induced with tunicamycin at 5 µg/mL for 4 hours. \**P* < 0.05; \*\**P* < 0.01; \*\*\**P* < 0.001; \*\*\*\**P* < 0.0001.

Exposure of cells to 4 hours of ER stress resulted in substantial changes in translation (**Fig. 3C, Supp. Fig. 2J**). Particularly, in agreement with previous work, we observed downregulation of translation for multiple transcripts and elevated ribosome occupancy of *ATF4* mRNA (**Supp. Fig. 2I**) ^18-22^. To address whether IGF2BP3 regulates the translation of UPR target genes, we analyzed the changes in the ribosome occupancy of translationally up- or downregulated UPR transcripts upon IGF2BP3 depletion (**Fig. 3C, D, Supp. Fig. 3J, K**). This analysis revealed that IGF2BP3 depletion reduced ribosome occupancy on translationally upregulated UPR targets bound by IGF2BP3. At the same time, IGF2BP3 depletion also reduced UPR-mediated translational repression, supporting a role for IGF2BP3 in eliciting a potent UPR.

### IGF2BP3 shapes the UPR through transcriptional feedback loops

Our genome-wide approaches revealed that IGF2BP3 depletion changed the levels of ∼15% of its target mRNAs identified by IR-PAR-CLIP, indicating that not all IGF2BP3-bound transcripts are affected by depletion of IGF2BP3 (**Supp. Fig. 3A**). Moreover, as mentioned above, we also found that the levels of UPR targets that are not directly bound to IGF2BP3 also decreased upon IGF2BP3 depletion, suggesting a secondary indirect effect of IGF2BP3 depletion (**Supp. Fig. 2C, D, H, J, K**). As IGF2BP3 regulates several mRNAs involved in regulation of transcription, we hypothesized that transcriptional reprogramming might result in the indirect effects observed upon IGF2BP3 depletion.

To uncover the primary mechanism of IGF2BP3-mediated regulation during ER stress, we aimed to uncouple transcriptional reprogramming from post-transcriptional regulation. To achieve this, we engineered a homozygous cell line expressing IGF2BP3 endogenously tagged with a mini auxin-inducible degron (mAID) in HCT116 cells stably expressing the OsTIR1 E3 ligase. This setup allowed for faster IGF2BP3 depletion than by CRISPR- or siRNA-based approaches, thus capturing mostly direct effects of IGF2BP3 loss. Auxin treatment depleted 95 % of mAID-IGF2BP3 in nine hours (**Supp. Fig. 3B, C**). Using these cell lines, we performed transcriptomics analyses following IGF2BP3 depletion in the absence and presence of ER stress (**Supp. Table 3**). The acute depletion of mAID-IGF2BP3 via auxin treatment increased levels of UPR target transcripts that are IGF2BP3-bound (**Fig. 4A**). Importantly, we observed a similar effect for all the IGF2BP3-bound transcripts, indicating a general destabilization effect of IGF2BP3 during ER stress (**Fig. 4A**). These data revealed that during ER stress, IGF2BP3 binding results in destabilization of IGF2BP3-bound UPR targets, supporting the hypothesis that a secondary indirect mechanism causes the observed dampening of the UPR.

**Figure 4.**
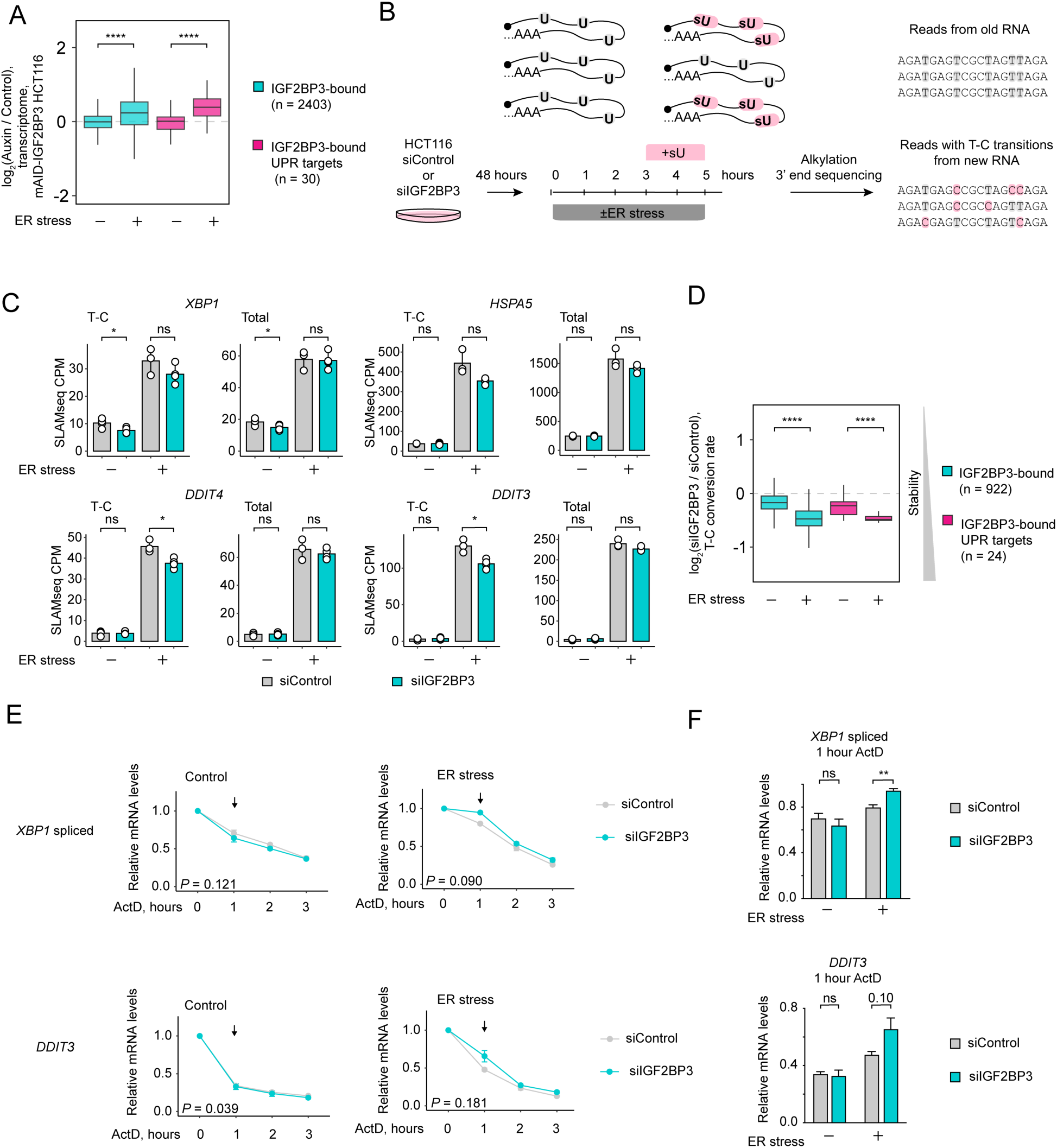
IGF2BP3 shapes the UPR through transcriptional feedback loops. **A.** Boxplot showing changes in transcript levels upon 9 hr auxin-induced mAID-IGF2BP3 depletion for IGF2BP3-bound transcripts and IGF2BP3-bound UPR targets. *P* values were calculated by two-sided Wilcoxon test. **B.** Design of thiol (SH)-linked alkylation for the metabolic sequencing of RNA (SLAMseq) experiment. IGF2BP3 is depleted for 48 hours using siRNA. For the last 5 hours before collection cells were treated with tunicamycin at 5 µg/mL or with DMSO as a control. In the last 2 hours 100 µM 4-thiouridine (s^4^U) was added to the cells to label the newly synthesized mRNA. **C.** Barplots showing SLAMseq total and T-C CPM values for selected UPR target genes (*HSPA5*, *XBP1*, *DDIT4*, and *DDIT3*). Values are the mean ± s.d of n=4 biological replicates. *P* values were calculated by two-sided Student’s t-test. **D.** Boxplot showing changes in estimated transcript stability T-C upon siRNA-mediated IGF2BP3 depletion for IGF2BP3-bound transcripts and IGF2BP3-bound UPR targets. The changes in transcript stability were assessed by analyzing the differences in average T-C conversion rates per gene following IGF2BP3 depletion. T-C conversion rate values indicate the relative abundance of both pre-existing and newly synthesized mRNA for a given gene, with a decrease in these values suggesting increased stabilization. *P* values were calculated by two-sided Wilcoxon test. **E.** RT-qPCR analyses of degradation rates for UPR target genes *XBP1* (spliced) and *DDIT3* upon siRNA-mediated depletion of IGF2BP3. HCT116 cells were treated with tunicamycin at 0.25 µg/mL for 4 hours and the transcription was blocked with 5 µg/mL ActD for indicated timepoints. Values are the mean ± s.e. of n=4 biological replicates. *P* values were calculated by paired two-sided Student’s t-test. **F.** Barplots showing total mRNA levels in unstressed cells and cells treated with tunicamycin at 0.25 µg/mL for 4 hours. Arrows indicate the 1-hour ActD treatment timepoint selected for boxplot representation in **E.** *P* values were calculated by two-sided Student’s t-test. \**P* < 0.05; \*\**P* < 0.01; \*\*\**P* < 0.001; \*\*\*\**P* < 0.0001.

To unravel whether transcriptional reprogramming contributes to the changes in the transcript levels upon IGF2BP3 depletion, we used thiol (SH)-linked alkylation for the metabolic sequencing of RNA (SLAMseq) method (**Fig. 4B**). In SLAMseq, nascent transcripts are metabolically labelled allowing assessment of *de novo* RNA synthesis ^50^. We performed two hours of pulse labeling in HCT116 cells with s^4^U, followed by total RNA extraction, alkylation, and mRNA 3′-end library preparation under homeostatic conditions and after subjecting cells to 5 hours of ER stress (with s^4^U added for the last two hours). We compared the control cells with the siRNA-based IGF2BP3-depletion to decipher the contribution of both post-transcriptional and transcriptional mechanisms driven by IGF2BP3 depletion (**Supp. Table 3**). Surprisingly, SLAMseq analyses showed that IGF2BP3 depletion decreased transcription of the UPR target genes (**Supp. Fig. 3D**), potentially due to downregulation of a transcriptional regulator upon IGF2BP3 depletion. Indeed, several mRNAs encoding for transcription factors were downregulated upon IGF2BP3 depletion (**Supp. Fig. 3E**). Comparison of *de novo* mRNA synthesis by SLAMseq with total mRNA levels measured in the same samples indicated that while IGF2BP3-bound UPR effectors *XBP1, HSPA5, DDIT4, and DDIT3* were less transcribed upon IGF2BP3 depletion this had a minor effect on total mRNA levels (**Fig. 4C**). This suggested that IGF2BP3 may have opposing transcriptional and post-transcriptional effects on the UPR target transcripts.

To systematically address the contribution of post-transcriptional regulation to the levels of the UPR target transcripts, we calculated the changes in mRNA stability by comparing the changes in the average T-C conversion rates per gene upon IGF2BP3 depletion (**Fig. 4D**). T-C conversion rate values reflect the relative abundance of pre-existing and *de novo* synthesized mRNA per given gene, and their decrease suggests stabilization. Those analyses showed that IGF2BP3 depletion increased the stability of mRNAs encoding for the UPR target genes, particularly during ER stress. These data indicated that IGF2BP3 depletion controls UPR transcript levels at the post-transcriptional and transcriptional levels.

Supporting the notion that IGF2BP3 destabilizes its target mRNAs during stress, when transcription was inhibited by actinomycin D (ActD) treatment, IGF2BP3 depletion increased the levels of *XBP1* and *DDIT3* (CHOP) mRNAs during ER stress (**Fig. 4E, F**). Notably, the destabilization effect was specific to ER stress but not to homeostatic conditions. These data suggested that IGF2BP3 regulates the UPR in two ways: 1. It binds to and destabilizes its target mRNAs during ER stress. 2. It indirectly upregulates the transcription of the UPR target genes. This transcriptional feedback loop allows differential regulation of UPR target transcripts by IGF2BP3 during ER stress. The transcriptional response is more potent, and this way, IGF2BP3 depletion dampens the response. This model explains how IGF2BP3 depletion leads to decreased levels of the UPR target transcripts while its depletion increases the levels of its post-transcriptional targets. It is attractive to speculate that the IGF2BP3-driven dual regulatory mechanism allows cells to decrease protein folding burden while ensuring they elicit potent UPR during ER stress.

### IGF2BP3 facilitates mRNA degradation during ER stress

Our transcriptomics and SLAMseq analyses showed that IGF2BP3-binding destabilizes most of its target mRNAs during ER stress (**Fig. 2D, 4A, D**). Notably, this was not observed in IGF2BP3 CRISPR/Cas9 knockout cells, where most of the genes showing the greatest differential expression decreased upon IGF2BP3 depletion in homeostatic conditions (**Fig. 2C**) and were unaffected by ER stress. It is plausible that this difference is due to cellular adaption to long-term IGF2BP3 depletion via transcriptional rewiring. Faster siRNA-mediated depletion resulted in both stabilization and destabilization of IGF2BP3-bound transcripts under homeostatic conditions, confirming that it regulates *HMGA2* (stabilized by IGF2BP3) ^44^ and *ZFP36L1* (destabilized by IGF2BP3) ^45^, as reported in earlier studies. Most top-regulated transcripts encoded proteins involved in cell proliferation and motility (e.g., *HMGA2*, *CARM1, TRIB1*, *NACC2*) ^51-54^. Notably, several of these target mRNAs encoded transcription factors (*ZNF385A*), regulators of gene expression (*HMGA2*, *CARM1, NACC2*), or RNA metabolism (*AGO2, ZFP36L1*), underlining that IGF2BP3-mediated posttranscriptional regulation impinges on potent gene regulatory networks to remodel the transcriptome.

The observed increase in IGF2BP3-mediated destabilization upon ER stress was specific for IGF2BP3-bound transcripts (**Fig. 5A, B**). These results were further confirmed using mAID-mediated IGF2BP3 depletion (**Supp. Fig. 4A**). In line with these analyses, a higher number of transcripts were destabilized by IGF2BP3 during ER stress compared to homeostatic conditions **(Fig. 5C, D**). Importantly, compared to homeostatic conditions, we also observed less potent IGF2BP3-mediated stabilization during ER stress **(Fig. 5C, D**). One of the most notable examples of ER-stress-modulated IGF2BP3 regulation was the well-described IGF2BP3 target mRNA *HMGA2*. *HMGA2* encodes a nonhistone chromatin factor that controls gene expression by altering chromatin architecture ^55^. In line with the published work, IGF2BP3 depletion led to lower levels of *HMGA2* mRNA under homeostatic conditions (**Fig. 5C, D, Supp. Fig. 4B, C**). The SLAMseq analyses confirmed that IGF2BP3 binding stabilizes *HMGA2* under those conditions (**Supp. Fig. 4D**). Surprisingly, *HMGA2* mRNA levels significantly decreased during ER stress (**Supp. Fig. 4B, C**). We confirmed that IGF2BP3 did not stabilize *HMGA2* mRNA as efficiently during ER stress as homeostatic conditions (**Fig. 5D, Supp. Fig. 4B, C, D**). These results were confirmed by actinomycin D chase experiments, which showed that IGF2BP3 depletion decreased the half-life of *HMGA2* mRNA under homeostatic conditions while not impacting it during ER stress (**Fig. 5E**). Notably, our IR-PAR-CLIP analyses showed that IGF2BP3 binding to *HMGA2* mRNA weakens during ER stress, which might explain the reduced protection of this RNA (**Fig. 1D**).

**Figure 5.**
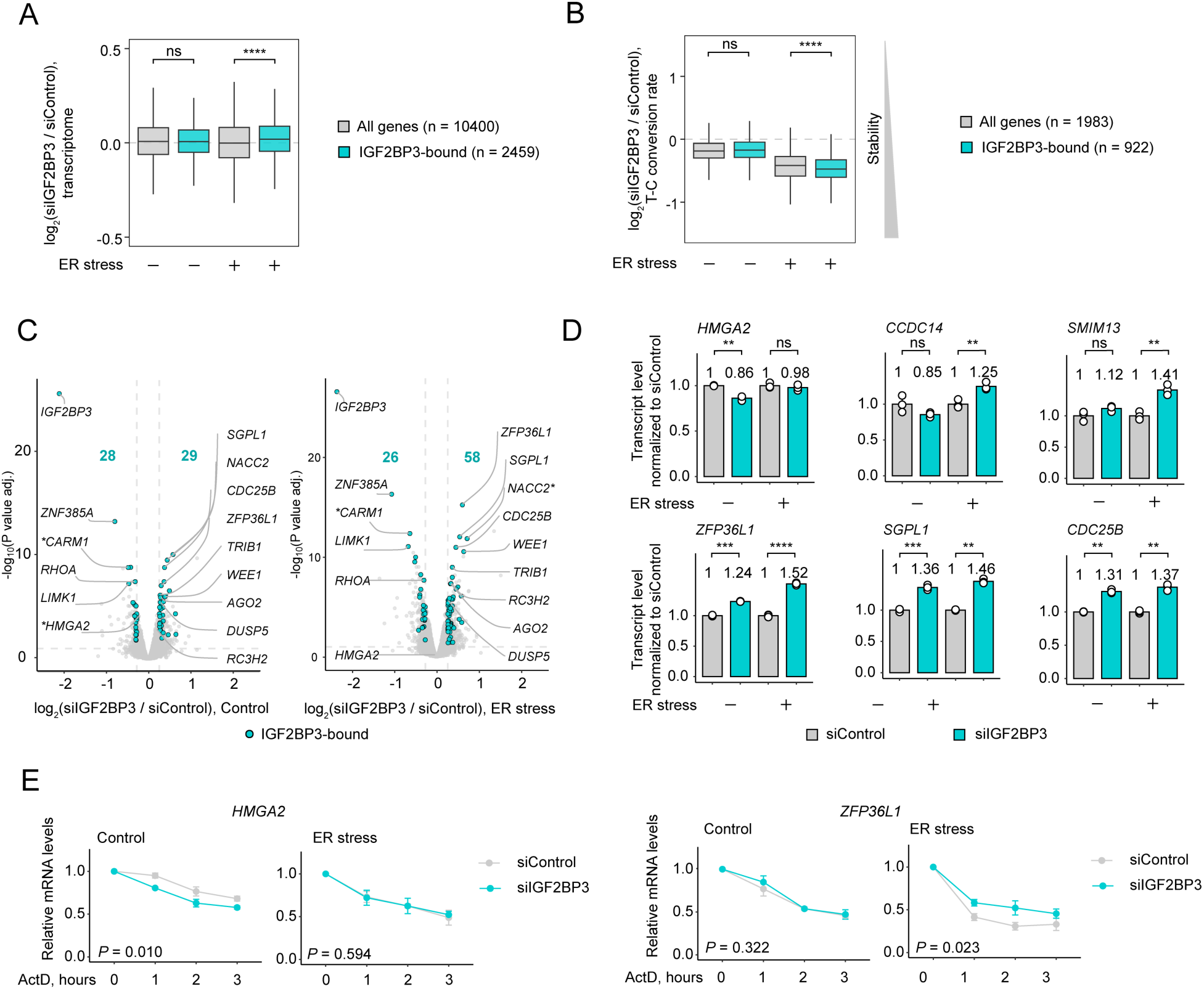
IGF2BP3 facilitates the degradation of its target mRNAs during ER stress. **A.** Boxplot showing changes in total transcript levels upon siRNA-mediated depletion of IGF2BP3 for all genes and IGF2BP3-bound transcripts in control and ER stress conditions. **B.** Boxplot showing changes in estimated transcript stability upon siRNA-mediated IGF2BP3 depletion for all genes and IGF2BP3-bound transcripts in control and ER stress conditions. The changes in transcript stability were assessed by analyzing the differences in average T-C conversion rates per gene following IGF2BP3 depletion. T-C conversion rate values indicate the relative abundance of both pre-existing and newly synthesized mRNA for a given gene, with a decrease in these values suggesting increased stabilization. *P* values were calculated by two-sided Wilcoxon test. **C.** Volcano plots of total transcriptome changes (RNA-seq) upon siRNA-mediated IGF2BP3 depletion with regulated genes highlighted (ΔRNA-seq > 20% and *P* value adj. < 0.05). * marks the genes encoding transcriptional regulators. **D.** Barplots showing total RNA-seq CPM values normalized to siControl conditions for selected genes regulated by IGF2BP3. Values are the mean ± s.d of n=3 biological replicates. *P* values were calculated by paired two-sided Student’s t-test. **E**. RT-qPCR analyses of degradation rates for *HMGA2* and *ZFP36L1* upon siRNA-mediated depletion of IGF2BP3. HCT116 cells were treated with tunicamycin at 5 µg/mL for 3 hours and the transcription was blocked with ActD for indicated timepoints. Values are the mean ± s.e. of n=3 biological replicates. *P* values were calculated by paired two-sided Student’s t-test. \**P* < 0.05; \*\**P* < 0.01; \*\*\**P* < 0.001; \*\*\*\**P* < 0.0001.

Consistent with these data, which suggest IGF2BP3 supports degradation of its targets during ER stress, ER stress resulted in more prominent destabilization for a specific subset of transcripts that IGF2BP3 also destabilizes under homeostatic conditions (**Fig. 5C, D, Supp. Fig. 4D**). IGF2BP3 depletion increased the levels of a well-described target, *ZFP36L1* mRNA under homeostatic conditions, supporting previous work ^45^ suggesting that IGF2BP3 mediates *ZFP36L1* mRNA degradation (**Fig. 5C, D, Supp. Fig. 4B, C, D**). This effect was more pronounced during ER stress, with *ZFP36L1* mRNA showing an increased half-life upon IGF2BP3 depletion under stress conditions (**Fig. 5D, E, Supp. Fig. 4C, D**). These analyses indicated that ER stress modulates IGF2BP3 function. Altogether, we discovered that ER stress modulates IGF2BP3-target mRNA interactions and impacts the regulation of its canonical targets.

### IGF2BP3 interacts with the mRNA degradation machinery

Association of IGF2BP3 with the mRNA degradation machinery was previously proposed to regulate a subset of its target RNAs ^45, 46, 56^. Therefore, we used proteomics to investigate whether changes in the IGF2BP3 interactome explain the increase in IGF2BP3-mediated degradation during ER stress. To this end, we immunoprecipitated IGF2BP3 from HCT116 cells. We also treated the samples with RNases to decipher protein-protein interactions mediated by IGF2BP3. The proteomics analyses showed that IGF2BP3 interacted with several proteins involved in RNA export (THO Complex subunits, MAGOH, RBM8A) and translation (ribosomal proteins) (**Supp. Fig. 5A, Suppl. Table 4**), supporting the model that IGF2BPs are loaded onto nascent transcripts in the nucleus and shuttle to the cytoplasm ^57^. IGF2BP3 also interacted with signal recognition particle (SRP) components, showing that IGF2BP3 binds to mRNAs during active translation en route to the ER, which is in line with our data.

Apart from those factors, our analyses identified association of IGF2BP3 with the RNA degradation machinery (RNA deadenylation and decapping machinery, exosome, XRN1) under both homeostatic conditions and during ER stress (**Fig. 6A, Suppl. Table 4**). These data supported the notion that IGF2BP3 accompanies its mRNA targets from synthesis to degradation (**Supp. Fig. 5A**). Several IGF2BP3-interacting proteins, including RNases IRE1 and XRN1 and components of the decapping machinery (EDC3, EDC4, DCP1A, DCP2) were insensitive to RNase treatment indicating protein-protein interactions (**Fig. 6B**).

**Figure 6.**
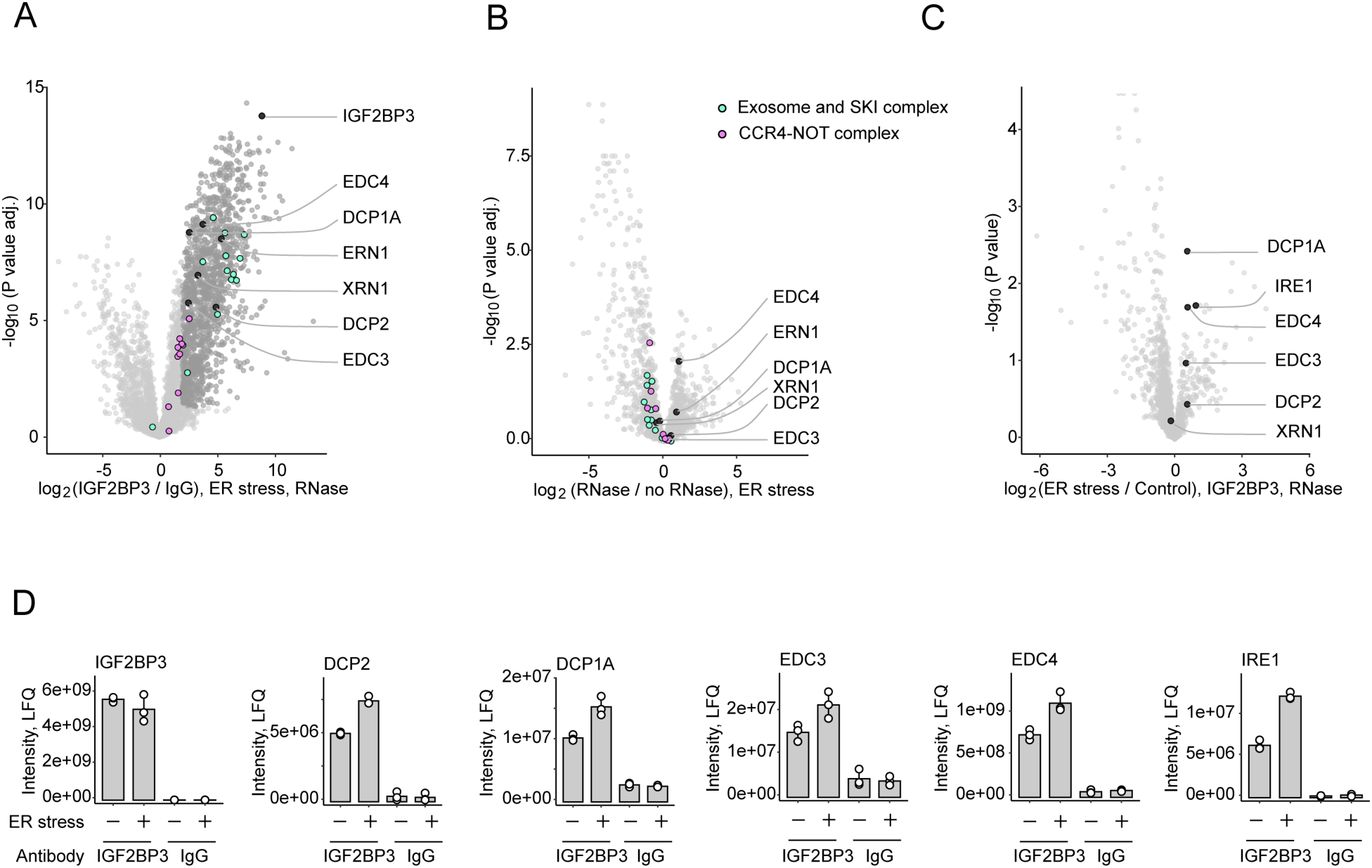
IGF2BP3 interacts with RNA degradation machinery. **A.** Volcano plot showing co-IP-MS analyses of endogenous IGF2BP3 (IGF2BP3 antibody over IgG control). IGF2BP3 interaction partners (4 times enriched over IgG control, *P* value adj. < 0.05) are shown in dark gray. **B.** Volcano plots comparing IGF2BP3 interactome with and without RNase treatment in ER stress conditions with highlighted functional groups of proteins. **C.** Volcano plots comparing IGF2BP3 interactome in control and ER stress conditions (tunicamycin at 5 µg/mL for 4 hours). **D.** Normalized LFQ values for selected examples of IGF2BP3 interacting proteins that increase interaction with IGF2BP3 upon ER stress. Values are the mean ± s.d. of n=3 biological replicates.

**Figure 7.**
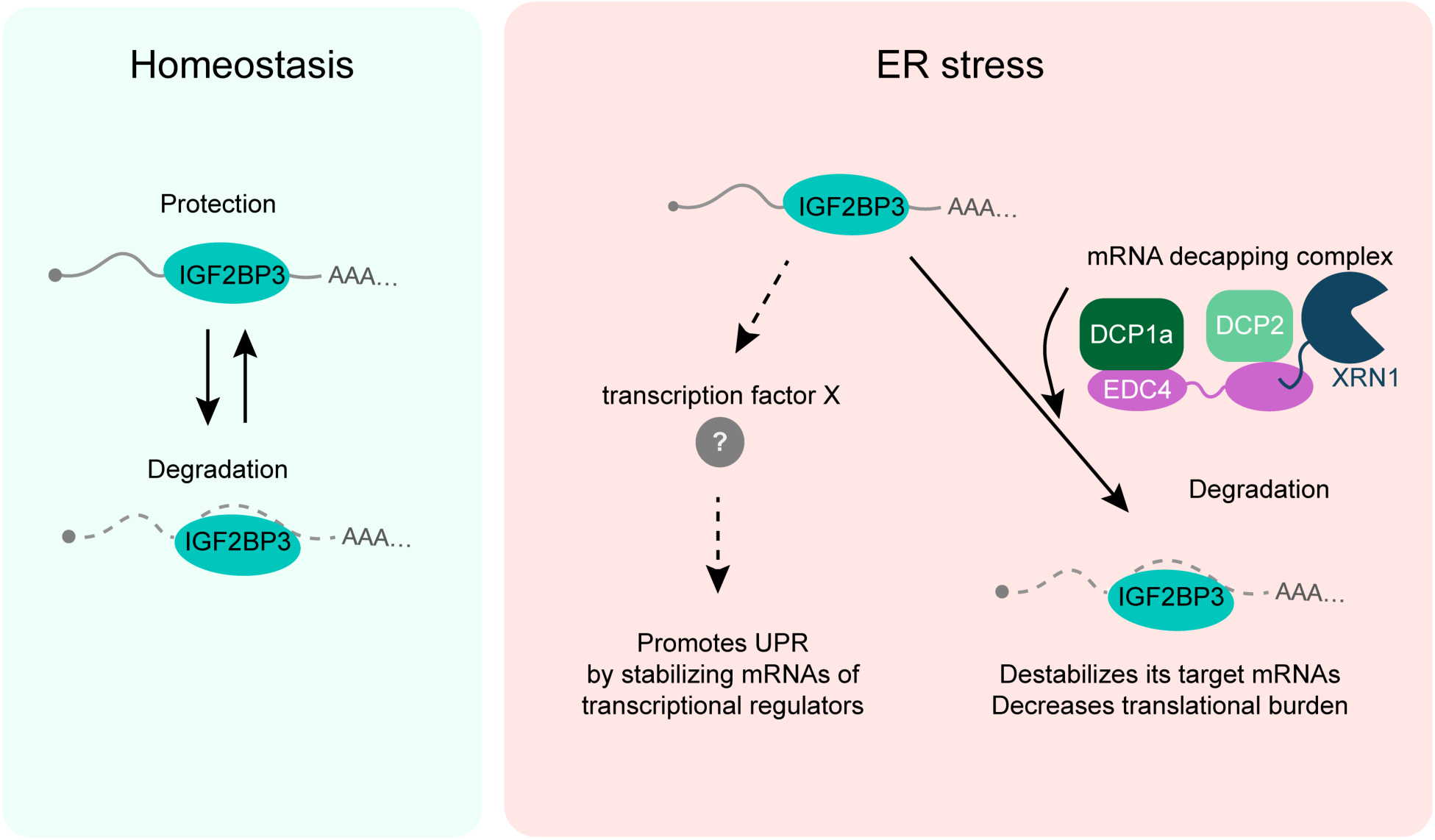
Model of IGF2BP3 function and regulation in ER stress conditions. Depending on the transcript identity IGF2BP3 binding to its’ target mRNAs can either promote their degradation or increase stability. Under homeostatic conditions there is an equal balance between stabilization and degradation by IGF2BP3. ER stress increases pro-degradation function of IGF2BP3 and promotes degradation of its target transcripts including the UPR effector mRNAs probably through increased interaction of IGF2BP3 with mRNA decapping complexes and ER stress sensor RNase IRE1. Upon ER stress, IGF2BP3 contributes to eliciting potent UPR indirectly through regulation of a yet unidentified transcription factor. This dual-layered regulation allows IGF2BP3 to elevate levels the levels of stress response gene expressions while decreasing other transcripts during ER stress to relieve protein-folding load.

While ER stress did not change IGF2BP3 interaction with most of these proteins, binding to a subset of proteins, including IRE1, increased, validating our initial findings (**Fig. 6C, D)**. Notably, in addition to IRE1, IGF2BP3 bound more strongly to all components of the mRNA decapping machinery (EDC3, EDC4, DCP1A, DCP2) during ER stress (**Fig. 6C, D**). This contrasted with XRN1, which bound to IGF2BP3 at similar levels in both conditions. These data suggested that increased interaction of IGF2BP3 with the mRNA decapping machinery might enable more efficient degradation of IGF2BP3 target mRNAs during ER stress. To summarize, our findings converge on the model that changes in the IGF2BP3 interactome shift its function towards promoting target transcript degradation during ER stress.

## Discussion

Among its many essential functions, the ER is the site for folding and maturation of secreted and transmembrane proteins, which form approximately one-third of the proteome. Cells need to rapidly adapt to changes in ER protein folding demands, as the separation of protein synthesis and folding in two distinct compartments poses a challenge. To overcome this, cells use diverse posttranscriptional and translational mechanisms to adjust the protein folding load in the ER. Here we discovered that IGF2BP3-driven posttranscriptional mechanisms facilitate mRNA degradation, as well as, indirectly tune transcription to generate gene regulatory networks that control UPR signaling while relieving protein-folding load on the ER.

Based on our published proteomics data on the interaction of the ER stress sensor IRE1 with IGF2BP3, here we explored the potential role of IGF2BP3 in posttranscriptional regulation of mRNA fate during ER stress. Our IR-PAR-CLIP analyses supported a model where IGF2BP3 predominantly binds to UPR target transcripts during ER stress, including those encoding the three master UPR transcription factors, XBP1, ATF4, and CHOP. Using complementary approaches to probe for posttranscriptional, transcriptional (RNA-Seq and SLAMseq), and translational changes (Ribo-Seq) upon IGF2BP3 depletion, we discovered a dual regulatory role mediated by IGF2BP3-driven posttranscriptional control that maintains cellular homeostasis during ER stress. While IGF2BP3 destabilizes most of its target mRNAs under ER stress conditions, it stabilized a subset of its target mRNAs, including key transcriptional regulators. Notably, the SLAMseq analyses revealed that an IGF2BP3-driven transcriptional feedback loop indirectly leads to the upregulation of many UPR genes. This IGF2BP3-driven dual regulation allows cells to downregulate the expression of most of the IGF3BP3 target transcripts while upregulating UPR-targets essential for adaptation to ER stress. This helps cells to decrease cellular folding burden while enabling them to efficiently express UPR target mRNAs. While the transcription factor that drives the IGF2BP3-driven feedback mechanism is currently unknown, multiple transcriptional regulators are sensitive to IGF2BP3 depletion. Several UPR transcription factors, including XBP1s and ATF6N, are bZIP transcription factors that form heterodimers with each other to regulate UPR combinatorially ^58^, it is plausible that some of the IGF2BP3-regulated bZIP transcription factors such as FOSL2 or ATF6 paralogue CREB3L2 might contribute to this effect.

In addition to IGF2BP3-mediated regulation of UPR effector RNAs, our data revealed that ER stress leads to a functional switch in IGF2BP3, increasing its destabilizing function. During heat shock stress, IGF2BP3 stabilizes its target mRNAs ^36^, suggesting that IGF2BP3 can drive opposing outcomes for its target transcripts depending on the cellular input. This dynamic and plastic response aligns with the cellular need to adjust transcript levels rapidly during proteotoxic stress. In line with these observations, we discovered that ER stress modulates IGF2BP3 interaction with its canonical targets. IGF2BP3 stabilizes *HMGA2* mRNA by protecting it from miRNA-driven degradation ^44^. During ER stress, IGF2BP3 binding to *HMGA2* mRNA decreased, resulting in less efficient stabilization of *HMGA2* transcript, and notably, this correlated with lower levels of *HMGA2*. HMGA2 regulates many biological processes involved in embryonic development, stem cell maintenance, and tumorigenesis ^55, 59, 60^. The resulting lower levels of HMGA2 during ER stress might interfere with early developmental processes, indicating a broader impact of stress-dependent regulation by IGF2BP3.

Our IP-MS/MS analyses of IGF2BP3 revealed that IGF2BP3 strongly associates with all the components of the mRNA decapping complex, an interaction which increased around two-fold during ER stress. We speculate that the increased association of IGF2BP3 with the mRNA decapping complex results in IGF2BP3 enhancing degradation during ER stress. Future work will provide insights into the mechanisms that mediate preferred association of the mRNA decapping complex with IGF2BP3 during ER stress.

IGF2BP3 is expressed at high levels in various tumors and its expression correlates with poor prognosis and cancer aggressiveness ^35, 46, 61^. ER stress is a prevalent feature of various cancers ^62^. Based on the metabolic state or cell identity, we speculate that the balance between IGF2BP3’s opposing outputs on UPR targets may shift, impacting UPR-driven cell fate decisions in cancer. Overall, our analysis of IGF2BP3’s association with target transcripts reveals a novel dual regulatory mechanism that ensures a robust response to ER stress.

## Acknowledgments

We are grateful to Niko Popitsch for support with the transcriptomics experiments and analyses of sequencing data. We are thankful to the Vienna BioCenter Next-Generation Sequencing Facility for deep sequencing services and Max Perutz Labs Proteomics Facility for mass spectrometry analyses. We thank Roland Foisner and Nana Naetar-Kerenyi for their input on CRISPR/Cas9 editing. RKO cells were a kind gift of the Johannes Züber lab. G.E.K. acknowledges funding from the Austrian Science Fund (FWF-SFB F79) and the Vienna Science and Technology Fund (WWTF-LS21-009). This research was partly funded by the Austrian Science Fund (FWF) [FWF-SFB F79]. For open access purposes, the author has applied a CC-BY public copyright license to any author-accepted manuscript version arising from this submission. A.S.A. is supported by the DOC Fellowship Programme of the Austrian Academy of Sciences.

## Materials and methods

### Mammalian cell culture

Mouse Embryonic Fibroblasts (MEF) expressing doxycycline-inducible GFP-IRE1 ^63^ were established in previous work ^64^. HCT116 WT cells were a kind gift from Prof. Manuela Baccarini (Max Perutz Labs). HCT116 cells conditionally expressing doxycycline-inducible (tetON) OsTIR1 were obtained from the Masato Kanemaki lab ^65^. RKO cells expressing doxycycline-inducible Cas9 were a kind gift from Johannes Zuber (IMP, Vienna, Austria) ^66^. HEK293T cells expressing splitGFP-IGF2BP3 were a kind gift of Manuel Leonetti established using published protocols ^67^.

HCT116 cells were cultured in McCoy’s 5A (modified) medium (Sigma, M9309) supplemented with 10% fetal bovine serum (Gibco, 10437028), 2 mM L-Glutamine (Sigma, G7513), 1% Pen/Step (Sigma, P0781). MEF cells were cultured in high-glucose DMEM media (Sigma, D5796) supplemented as above. RKO cells were cultured in RPMI-1640 media (Sigma, R8758) supplemented with 10% fetal bovine serum (Gibco, 10437028), 2 mM L-Glutamine (Sigma, G7513), 1% Pen/Step (Sigma, P0781), 1x non-essential amino acids (Thermo Scientific, 11140050), and 2 mM sodium pyruvate (Gibco, 11360070). All cell lines were cultured in a humidified incubator at 37°C and 5% CO2 and regularly tested for Mycoplasma infection with the EZ-PCR™ Mycoplasma Detection Kit (Biological Industries).

### Western blotting

80% confluent cells were lysed with RIPA buffer (150 mM NaCl, 1% NP-40, 0.5% Sodium deoxycholate, 0.1% SDS, and 25 mM TRIS pH 7.4) with 1x EDTA-free protease inhibitor cocktail (Roche). Lysates were clarified using a table-top centrifuge at maximum speed (20,000 g) for 20 min at 4°C. Western blot samples were denatured at 95 °C for 5 min in 1x SDS sample buffer (50 mM Tris-HCl pH 6.8, 2% SDS, 0.1% Bromophenol blue, 10% glycerol, 20 mM DTT). Following denaturing, the samples were loaded onto a 10%, 12% or 15% sodium dodecyl sulfate (SDS) gel. Proteins were transferred onto a 0.2 µm nitrocellulose membrane (Amersham) using BioRad Trans-Blot Turbo Transfer System or with wet transfer in transfer buffer (25 mM TRIS, 190 mM glycine, 20% ethanol) for 110 min at 120 V. Membranes were stained with Ponceau S and blocked in 5% milk for 1 hour. The primary antibody (**Supp. Table 5**) was diluted in 2.5% milk and incubated overnight at 4 °C. The membrane was washed 5 times with TBST (20 mM TRIS, 150 mM NaCl, 0.1% Tween 20), and the membranes were incubated for 1 h with the secondary antibody (**Supp. Table 5**) diluted in 2.5% milk. After the incubation the membranes were washed 5 times with TBST. Membranes were developed with enhanced chemiluminescent (ECL) horse radish peroxidase substrate (WESTAR ETA C ULTRA 2.0, Cyanagen), imaged using BioRad ChemiDoc, and analyzed using ImageLab software (6.1.0).

### Co-Immunoprecipitation of GFP-IRE1

Two ø15 cm dishes of 80% confluent MEF GFP-IRE1 cells were used per condition. To induce ER stress, cells were treated with 5 µg/mL tunicamycin (TM) for 4 hours (if other time is not indicated), DMSO in 1:1000 dilution was used a control. Cells were washed with ice-cold PBS and collected by scraping and pelleting at 500 g at 4°C for 5 min. Cells were lysed in 250 µL of lysis buffer (25 mM HEPES pH 7.3, 150 mM NaCl, 1% NP-40, 1 mM EDTA, 10% Glycerol, 1x EDTA-free protease inhibitor cocktail, 0.4 U/µL RNasin RNase inhibitor) per condition by incubation on ice 10 min with intermittent vortexing. The lysate was clarified with two steps of centrifugation at 4°C 1,000 g for 5 min and 13,000 g for 15 min. The immunoprecipitation (IP) was performed with GFP-trap magnetic beads (ChromoTek). 40 µL of bead slurry was used for two ø15 cm dishes. The lysate was incubated with the GFP-trap beads for 2 hours at 4°C and the beads were washed 3 times with 1 mL of ice-cold lysis buffer. For RNase digestion 0.1 U/µL of RNase I (Ambion) and 1 U/µL of RNase T1 (EN0541) were added to the second wash. For the control (undigested) samples 1 U/µL of RNasin RNase inhibitor was added. Samples were incubated at room temperature for 10 min on a rotator. Proteins were eluted in 25 µL of 1x SDS sample buffer at +95 °C for 5 min, loaded on SDS-PAGE and analyzed by western blotting as described above with anti-IGF2BP3, anti-GFP and anti-GAPDH antibodies (**Supp. Table 5**). For the GFP-IGF2BP3 co-IP from HEK293T same conditions were used as for MEF cells. One ø15 cm dish was used per condition; to induce ER stress, the cells were treated with 5 µg/mL TM for the indicated timelines or for 4 hours.

### IR-PAR-CLIP of IGF2BP3

#### IGF2BP3 IR-PAR-CLIP library preparation and data processing

The detailed protocol for the infrared (IR) PAR-CLIP of IGF2BP3 is described in (Anisimova, Karagöz, 2023). 125 million (5x ø15 cm dishes) HCT116 inducible tetON OsTIR1 cells were used per condition. 100 μM 4sU (Sigma) was added to the cell culture media 15 h prior to collection. ER stress was induced with 5 µg/mL TM for 4 hours prior to collection. DMSO in 1:1000 dilution was used a control. IGF2BP3 was immunoprecipitated with anti-IGF2BP3 Proteintech antibody (14642-1-AP, lot 00088732). anti-IgG Proteintech antibody (30000-0-AP) was used as a control. RNase I (Ambion) was used for the in-lysate RNase digestion at 0.1 U/µL and for the on-bead digestion in 1 mL of the lysis buffer at 0.025 U/µL. IR-PAR-CLIP libraries were sequenced on a NovaSeq 6000 S1 on SR100 (Illumina) at the Vienna BioCenter NGS facility with 55 million reads per sample on average. UMIs were extracted and adapter sequences were trimmed using UMI-tools v1.1.1 (Smith et al. 2017). The reads were size- and quality-trimmed using Trimmomatic v0.30 (Bolger et al. 2014) to have a length between 18 and 45 nt. The reads were then mapped to human genome hg38 using GENCODE annotation (release 36) with bowtie v0.12.7 (Langmead et al. 2009), allowing up to three mismatches and deduplicated using UMI-tools v1.1.1. Gene counts were obtained using the FeatureCounts function of the Subread package v2.0.1^68^. Gene counts for protein coding genes were RLE normalized to calculate the counts per million values (CPM), filtered to only include genes with CPM higher than 5 in at least one third of the libraries, and the differential expression analysis was performed with edgeR glmQLFTest (Generalized Linear Model Quasi-Likelihood F-test) ^69^.

### IR-PAR-CLIP matching transcriptome library preparation and data processing

An aliquot of the lysate was taken before the in-lysate RNase digest for total RNA isolation and sequencing. Total RNA was isolated using peqGOLD TriFast (Peqlab, VWR) and transcriptome libraries were prepared with QuantSeq 3’ mRNA-Seq Library Prep Kit (Lexogen). Libraries were sequenced on a NextSeq2000 P2 at SR100 mode (Illumina) at the Vienna BioCenter NGS facility with 30 million reads per sample on average. The quality-, adapter-, and polyA-trimmed reads were aligned to human genome hg38 using GENCODE annotation (release 36) with STAR v2.7.5c. As the reads originate from cross-linked PAR-CLIP samples and have T to C transitions, mismatches were allowed. Gene counts for protein coding genes were RLE normalized to calculate the counts per million values (CPM), filtered to only include genes with a CPM higher than 1 in at least half of the libraries. Differential expression analysis was performed with edgeR glmQLFTest ^70^.

### IR-PAR-CLIP computational analysis

To estimate relative binding of IGF2BP3, IR-PAR-CLIP CPMs were divided to the matching total transcriptome QuantSeq CPMs. A pseudocount of 1 CPM was added to all samples and only genes with mean CPM for QuantSeq samples greater than 5 CPM were taken for the analysis. To identify IGF2BP3 target genes, IGF2BP3-binding clusters were called with PARalyzer v1.5 ^71^ (ini file is available in **Supp. Table 5**). Genes that had at least one cluster containing more than 25 CPM and more than 50% of T to C conversions per read in two replicates (number 2 and 3) per condition were selected as IGF2BP3 targets. Samples from replicate 1 were excluded from target identification due to lower final read number and therefore lower number of identified clusters and targets. The target lists were additionally filtered to only include targets with PAR-CLIP CPMs higher than total transcriptome QuantSeq CPMs. PAR-CLIP genomic tracks were visualized using svist4get v1.2.20 ^72^. The enrichment of 5-nt motifs in IGF2BP3 PAR-CLIP reads was analyzed with HOMER v4.11 ^73^, findMotifs.pl command using shuffled background. Sequencing data processing was done using the HPC of the Center for Integrative Bioinformatics Vienna (CIBIV) and Life Science Compute Cluster (LiSC) of the University of Vienna, Austria.

### Establishment of IGF2BP3 knockout cell lines and siRNA knockdown

For IGF2BP3 knockout cell line generation, gRNA sequences (clones E4 and A3: 5 ′ TGGCACCGACTGATAGAGCT 3′; clone D12: 5′ACGCGTAGCCAGTCTTCACC 3′) were cloned into the pSpCas9 (BB)-2A-GFP (PX458) (plasmid #48138; Addgene) ^74^. HCT116 tetON OsTIR1 cells were transiently transfected using jetOPTIMUS reagent (Tamar, 101000051), and GFP-positive single-cell clones were FACS sorted at BD FACSAria IIIu at Max Perutz Labs BioOptics FACS Facility. For siRNA knockdown, HCT116 cells (WT or tetON OsTIR1) or RKO iCas9 were transfected with IGF2BP3 (Dharmacon, L-003976-00-0005) SMARTpool siRNAs using DharmaFECT 2 (Dharmacon, T-2002-01) at 75 nM for 48 hours. ON-TARGETplus nontargeting siRNA pool (Dharmacon, D-001810-10-05) was used as a control.

### Total transcriptome sequencing (RNA-Seq) of IGF2BP3 KO HCT116 cells

HCT116 tetON OsTIR1 IGF2BP3 KO clones E4, A3, and D12 (in duplicates) and parental cells (four replicates) were treated with 5 µg/mL TM for 4 hours (1:1000 DMSO was used as a control). Total RNA was isolated using peqGOLD TriFast (Peqlab, VWR), treated with RNase-free DNase I (NEB), and re-purified with peqGOLD TriFast (Peqlab, VWR). RNA precipitation was done using the isopropanol method. Total RNA sequencing libraries were prepared with NEBNext Poly(A) mRNA Magnetic Isolation Module (NEB, E7490L) and NEBNext Ultra II Directional RNA Library Prep Kit (NEB, E7765L) by Vienna BioCenter NGS facility sequenced on a NovaSeq S4 on PE100 (Illumina) with 55 million reads per sample on average. The quality- and adapter-trimmed reads were aligned to human genome hg38 using GENCODE annotation (release 36) with STAR v2.7.5c allowing up to two mismatches per read. Gene counts for protein coding genes were RLE normalized to calculate the counts per million values (CPM), filtered to only include genes with CPM higher than 5 in at least half of the libraries, and differential expression analysis was performed with edgeR glmQLFTest ^70^.

### Ribosome profiling (Ribo-Seq) and RNA-Seq of HCT116 cells upon siRNA-mediated depletion of IGF2BP3

HCT116 WT cells were depleted of IGF2BP3 using siRNA pools as described in “Establishment of IGF2BP3 knockout cell lines and siRNA knockdown” section. ER stress was induced with 5 µg/mL TM for 4 hours prior to collection. One ∼80% confluent well of a 6-well plate per condition (three biological replicates per condition) was washed with ice-cold PBS (Sigma) supplemented with 100 mg/mL cycloheximide, lysed on a plate on ice with 350 μL of ice-cold lysis buffer (20 mM HEPES pH 7.3, 150 mM KCl, 5 mM MgCl2, 1% Triton X-100, 100 mg/mL cycloheximide, 1 mM DTT, 1x EDTA-free protease inhibitor cocktail), scraped, and transferred to 1.5 mL RNase-free tube. Cell lysates were passed three times through the 27G needle, incubated on ice for 10 min with intermittent vortexing, and clarified on a table-top centrifuge at maximum speed (20,000 g) for 20 min at 4°C. Total RNA concentration in the lysate was estimated using OD A260 measurement on Nanodrop. The lysate concentrations were brought to 240 ng/μL, 15 μL aliquots were taken for matching RNA-Seq. 200 μL of the lysate were supplemented with 5 mM CaCl_2_, 2 U/μL DNase I (NEB), 0.65 μg of micrococcal nuclease (Sigma, 10107921001), and 0.7 U of RNase I (Ambion) and incubated at room temperature for one hour on a rotator. The lysate was layered on 10%-50% sucrose gradient and centrifugated at 35,000 g for 3 hours at 4°C in a SW40Ti rotor. Polysome fractions were separated on a BioComp fractionator and monosome fractions were collected. Ribo-Seq libraries were prepared as described in ^75^ with the following changes. rRNA was depleted from samples after adapter ligation using human riboPOOL probes (siTOOLs) according to the manufacturer’s instructions, purified with Oligo Clean & Concentrator kit, and eluted in 12 μL of RNase-free water. Reverse transcription was done with SuperScript III (Invitrogen). Oligonucleotides used for Ribo-Seq are listed in **Supp. Table 5**. Libraries were sequenced on a NovaSeq SP on SR100 (Illumina) at the Vienna BioCenter NGS facility with 60 million reads per sample on average. RNA-Seq libraries were prepared with NEBNext Poly(A) mRNA Magnetic Isolation Module (NEB, E7490L) and NEBNext Ultra II Directional RNA Library Prep Kit (NEB, E7765L) and sequenced on a NovaSeq SP on SR100 (Illumina) with 30 million reads per sample on average. The quality- and adapter-trimmed reads were aligned to human genome hg38 using GENCODE annotation (release 36) with STAR v2.7.5c allowing up to one or two mismatches per read for Ribo- and RNA-Seq respectively. Gene counts for protein coding genes were filtered to include genes with coding sequence length-normalized CPM values higher than 5 in at least one third of the libraries. Differential expression analysis was performed with edgeR glmQLFTest ^70^. For ribosome occupancy analysis changes in Ribo-Seq were compared to changes in RNA-Seq with edgeR glmQLFTest ^70^.

### Establishment of doxycycline-inducible IGF2BP3 knockout RKO cell lines

RKO-Dox-Cas9 (iCas9) cell lines for doxycycline-inducible knockout IGF2BP3 were established using lentiviral transduction with lentiviral particles (produced as described in ^76^ containing Dual-sgRNA_hU6-mU6 vectors described in ^66^ expressing two sgRNAs against IGF2BP3 (5 ′ TGGCACCGACTGATAGAGCT 3′ and 5′ GAAGATACTTTCTAGGTCCG 3′) or against non-coding locus AAVS1 (5’ CGCTGTGCCCCGATGCACAC 3’ and 5 ′ GGCGCGTCGCTCGCTCGCTC 3′) from human and mouse U6 promoters and eBFP2 from a PGK promoter. The eBFP2-positive cells were FACS sorted at BD FACSMelody™ Cell Sorter at Max Perutz Labs BioOptics FACS Facility.

### SLAMseq of RKO iCas9 and HCT116 cells upon IGF2BP3 depletion

IGF2BP3 depletion was induced in RKO iCas9 cells with 250 ng/mL doxycycline for 72 hours. siRNA-mediated depletion of IGF2BP3 was performed in either RKO iCas9 or HCT116 WT as described in the “Establishment of IGF2BP3 knockout cell lines and siRNA knockdown” section. ER stress was induced in HCT116 WT cells using a 5 hour-long treatment with 5 µg/mL TM (1:1000 DMSO was used as a control). 4-thiouridine (s^4^U) was added to cell culture media at 250 µM 2 hours prior collection. Total RNA was extracted using the KingFisher Flex Purification System (Thermo) with the High-Performance RNA Isolation kit (Molecular Tools Shop, Vienna BioCenter). During the isolation RNA was treated with RNase-free DNase I (NEB). SLAMseq samples were prepared according to the standard SLAMseq protocol described in ^50^. Briefly, total RNA was alkylated with 10 mM iodoacetamide in alkylation buffer (50 mM sodium phosphate buffer pH 8.0, 50% DMSO) at 50°C for 15 min and purified using the KingFisher Flex Purification System. Sequencing libraries were prepared with NEBNext Poly(A) mRNA Magnetic Isolation Module (NEB, E7490L) and NEBNext Ultra II Directional RNA Library Prep Kit (NEB, E7765L) and sequenced on a NovaSeq SP on SR100 (Illumina) at the Vienna BioCenter NGS facility with 30 million reads per sample on average. The quality- and adapter-trimmed reads were aligned to human genome hg38 using GENCODE annotation (release 36) with STAR v2.7.5c allowing up to five mismatches per read. Gene counts for protein coding genes were filtered to include genes with coding sequence length-normalized CPM > 5 in at least half of the libraries. Differential expression analysis was performed with edgeR glmQLFTest ^70^. The data were processed using the HPC of the Center for Integrative Bioinformatics Vienna (CIBIV).

### Identification of siRNA pool off-targets from siRNA knockdown experiments

To exclude possible siRNA off-target genes we compared changes in gene expression upon doxycycline-inducible knockout and siRNA-mediated depletion of IGF2BP3 in RKO iCas9 cells. Genes were considered off-targets if changes in their levels were more than 20% upon siRNA-mediated depletion, but less than 10% upon doxycycline-inducible knockout. Significance was not considered for this analysis due to overall small amplitude of changes. The identified off-targets are listed in **Supp. Table 5**. These genes were excluded from analyses of experiments where IGF2BP3 was depleted with siRNA pools.

### RNA isolation and RT-qPCR

Total RNA was isolated using the KingFisher Flex Purification System (Thermo) with the High-Performance RNA Isolation kit (Molecular Tools Shop, Vienna BioCenter). During the isolation RNA was treated with DNase I (M0303S, NEB). cDNA was prepared with LunaScript RT SuperMix (NEB) and amplified in a qPCR reaction with 2x Hot Start qPCR master mix (Molecular Tools Shop, Vienna BioCenter) using BioRad CFX 384 Touch machine. The qPCR primers are listed in **Supp. Table 5**. mRNA levels were calculated relative to *RPL6* levels.

### Generation of the HCT116 cell line expressing IGF2BP3 endogenously tagged with mini auxin inducible degron (mAID)

The HCT116 mAID-IGF2BP3 cell line was generated according to the protocol described in ^77^. To endogenously tag IGF2BP3 with mAID, sequences containing N-terminal homology arms (629 bp before and 78 bp after the Cas9 cut site) of IGF2BP3 with SacI, SalI, BamI, and XhoI restriction sites were ordered from IDT. The gene block was amplified using gene block adapter primers. The IGF2BP3 homology arms were inserted into the pMK344 plasmid backbone (Addgene #121179) using SacI and XhoI restriction sites. BSD-P2A-mAID sequence was cloned into the plasmid with N-terminal homology arms based on IGF2BP3 from the pMK347 plasmid (Addgene #121181) using SalI and BamI restriction sites resulting in the HDR template. The PAM sites were mutated in the HDR template using site-directed mutagenesis with a primer pair (IGF2BP3_PAM_mut_F and IGF2BP3_PAM_mut_R (**Supp. Table 5**)). To obtain the Hygro-P2A-mAID HDR template, Hygro-P2A-mAID was cloned from pMK344 to replace the BSD-P2A-mAID in the final IGF2BP3 HDR template with mutated PAM sites using SalI and BamI restriction sites. To ensure the homozygous insertion of mAID-IGF2BP3 into the genome, HCT116 tetON OsTIR1 cells ^78^ were transiently transfected with a 1:1:2 ratio mixture of the BSD-P2A-mAID HDR, Hygro-P2A-mAID HDR template plasmids and pSpCas9 (BB)-2A-GFP (PX458) (plasmid #48138, Addgene) ^74^ targeting the first exon of the IGF2BP3 gene (**Supp. Table 5**) using the Fugene HD (Promega) reagent according to the manufacturer’s instructions. 24 hours after transfection, cells were collected by trypsinization and plated at a 1:200 dilution in standard culture media (McCoy’s 5A (modified) with 10% FBS, 2 mM L-Glutamine, 1% Pen/Step). The next day, the media was supplemented with 50 µg/mL of Hygromycin B Gold (InvivoGen, #ant-hg) and 50 µg/mL of Blasticidin S Hydrochloride (InvivoGen, #ant-bl-05). Cells were grown in selection media until visible colonies were formed and expanded in 96-well plates. The homozygous insertion was verified using western blotting with anti-IGF2BP3 and anti-GAPDH antibodies and genotyped using DirectPCR Lysis-Reagent Cell (Peqlab, VWR) with IGF2BP3_CDS_1_R, IGF2BP3_5UTR_1_F, HygR_F, and BSDR_F primers. All primer sequences are listed in **Supp. Table 5**.

### SLAMseq of HCT116 tetON OsTIR1 cells upon auxin-mediated depletion of mAID-IGF2BP3

When expressed in mammalian cells OsTIR1 forms a complex with SCF (Skp1–Cul1–F box) components form E3 ubiquitin ligase that recognizes mAID upon addition of auxin (IAA) ^77^. To induce expression of OsTIR1, cells were treated with 200 ng/mL doxycycline for 24 hours. mAID-IGF2BP3 depletion was induced with IAA added to the cell culture media at 500 μM for 9 hours and ER stress was induced with 5 µg/mL TM for 5 hours (1:1000 DMSO was used as a control). 4-thiouridine (s^4^U) was added to cell culture media at 250 µM for 2 hours. All indicated treatment times are hours prior to collection. One ∼80% confluent well of 24-well plate was used per condition, and the experiment was performed in quadruplicate. RNA isolation, SLAMseq library preparation and analysis were performed as described above in the “SLAMseq of RKO iCas9 and HCT116 cells upon IGF2BP3 depletion” section.

### Co-Immunoprecipitation of endogenous IGF2BP3 followed by mass spectrometry

One ø15 cm dish of 80% confluent HCT116 WT cells was used per condition. To induce ER stress, cells were treated with 5 µg/mL tunicamycin (TM) for 4 hours, and DMSO at 1:1000 dilution was used as a control. Cells were washed with ice-cold PBS and collected by scraping and pelleting at 500 g at 4°C for 5 min. Cells were lysed in 250 µL of lysis buffer (25 mM HEPES pH 7.3, 150 mM NaCl, 0.5% NP-40, 0.5 mM EDTA, 10% Glycerol, 1x EDTA-free protease inhibitor cocktail, 1x PhosSTOP) per condition by incubation on ice 15 min with intermittent vortexing. The lysate was clarified by centrifugation at 20,000 g for 20 min at 4°C. 10 μg of Proteintech anti-IGF2BP3 antibody (14642-1-AP, lot 00088732) or Proteintech IgG control (30000-0-AP) was coupled to the 40 μL of protein G Dynabeads (Invitrogen) in 1 mL of lysis buffer for 20 min, rotating at room temperature, washed three times with 1 mL of the lysis buffer, resuspended in the original bead volume (40 μL) and added to 200 µL of the clarified lysate. Beads were washed five times with 1 mL of wash buffer (25 mM HEPES pH 7.3, 150 mM NaCl, 0.5 mM EDTA, 10% Glycerol). Each wash was incubated on ice for 3 min. For RNase digestion 0.1 U/µL of RNase I (Ambion) and 1 U/µL of RNase T1 (EN0541) were added to the second wash. Samples were incubated at room temperature for 10 min on a rotator. The control (undigested) samples were kept on ice. The proteins were eluted from the beads with 20 µL 100 mM glycine pH 2.0 three times and the pooled supernatant was adjusted to alkaline using about 20 µL 1M Tris pH 8.0. Disulfide bonds were reduced with 3.2 µL of 250 mM dithiothreitol (DTT) for 30 min at room temperature before adding 3.2 µL of 500 mM iodoacetamide and incubating for 30 min at room temperature in the dark. Remaining iodoacetamide was quenched with 1.6 µL of 250 mM DTT for 10 min. Proteins were digested with 300 ng trypsin (Trypsin Gold, Promega) in 3 µL 50 mM ammonium bicarbonate at 37°C overnight. The digest was stopped by the addition of 10% trifluoroacetic acid (TFA) to a final concentration of 0.5%. The peptides were desalted using C18 Stagetips ^79^.

### Liquid chromatography-mass spectrometry analysis

Peptides were separated on a Vanquish Neo nano-flow chromatography system (Thermo-Fisher), using a trap-elute method for sample loading (Acclaim PepMap C18, 2 cm × 0.1 mm, 5 μm, Thermo-Fisher), and a C18 analytical column (Acclaim PepMap C18, 50 cm × 0.075 mm, 2 μm, Thermo-Fisher), applying a segmented linear gradient from 2% to 35% and finally 80% solvent B (80 % acetonitrile, 0.1 % formic acid; solvent A 0.1 % formic acid) at a flow rate of 230 nL/min over 120min. Eluting peptides were analyzed on an Exploris 480 Orbitrap mass spectrometer (Thermo-Fisher Scientific) coupled to the column with a FAIMS pro ion-source (Thermo-Fisher Scientific) using coated emitter tips (PepSep, MSWil) with the following settings: The mass spectrometer was operated in DDA mode with two FAIMS compensation voltages (CV) set to −45 or −60 and 1.5 s cycle time per CV. The survey scans were obtained in a mass range of 350-1500 m/z, at a resolution of 60k at 200 m/z, and a normalized AGC target at 100%. The most intense ions were selected with an isolation width of 1.4 m/z, fragmented in the HCD cell at 30% collision energy, and the spectra recorded for max. 50 ms at a normalized AGC target of 100% and a resolution of 15k. Peptides with a charge of +2 to +6 were included for fragmentation, the peptide match feature was set to preferred, the exclude isotope feature was enabled, and selected precursors were dynamically excluded from repeated sampling for 45 seconds.

### Proteomics data analysis

The RAW MS data were analyzed with FragPipe (20.0), using MSFragger (4.1) ^80^. IonQuant (1.10.27) ^81^ and Philosopher (5.0.0) ^82^. The default FragPipe workflow for label free quantification (LFQ-MBR) was used, except “Normalize intensity across runs” was turned off. Cleavage specificity was set to Trypsin/P, with two missed cleavages allowed. The protein FDR was set to 1%. A mass of 57.02146 (carbamidomethyl) was used as fixed cysteine modification; methionine oxidation and protein N-terminal acetylation were specified as variable modifications. MS2 spectra were searched against the human 1 protein per gene reference proteome from Uniprot (Proteome ID: UP000005640, release 2024_01), concatenated with a database of 382 common laboratory contaminants (release 2023.03, https://github.com/maxperutzlabs-ms/perutz-ms-contaminants).

Computational analysis was performed using Python and the in-house developed Python library MsReport (version 0.0.24, ^83^). LFQ protein intensities reported by FragPipe were log2-transformed and normalized across samples using the ModeNormalizer from MsReport. The missing normalized LFQ intensity values were imputed by drawing random values from a normal distribution after filtering out contaminants, proteins with less than 2 peptides and less than 2 quantified values in at least one group. Differences between groups were statistically evaluated using the LIMMA 3.52.1 ^84^ at 5% FDR (Benjamini-Hochberg). The in-house Python library XlsxReport (0.1.0) was used to create a formatted Excel file summarizing the results of protein quantification. Proteins were considered IGF2BP3 interactors if their normalized LFQ values were enriched more than 4 times over IgG control, *P* value adj. < 0.05. GO term analysis was performed with ShinyGo webserver ^85^.

## Data deposition

Next generation sequencing data have been deposited in GEO with the following identifiers:

GSE289214 – RNA-Seq of HCT116 cells upon IGF2BP3 CRISPR/Cas9 knockout in control and ER stress conditions

GSE289023 – IR-PAR-CLIP of IGF2BP3 in HCT116 cells in control and ER stress conditions

GSE289024 – transcriptome (QuantSeq) of HCT116 cells in control and ER stress conditions matching IR-PAR-CLIP samples

GSE289425 – Ribo-Seq of HCT116 cells upon IGF2BP3 siRNA knockdown knockout in control and ER stress conditions

GSE289424 – transcriptome (RNA-Seq) of HCT116 cells upon IGF2BP3 siRNA knockdown knockout in control and ER stress conditions matching Ribo-Seq samples

GSE289482 – SLAMSeq of HCT116 cells upon IGF2BP3 knockdown in control and endoplasmic reticulum (ER) stress conditions

GSE289481 – SLAMseq of auxin-mediated depletion of mAID-IGF2BP3 in control and endoplasmic reticulum (ER) stress conditions. Depletion is induced by auxin (IAA) treatment in mAID-IGF2BP3 cells, parental (WT) cells are used as a control.

GSE289480 – SLAMseq of doxycycline-inducible knockout and siRNA-mediated depletion of IGF2BP3 in RKO iCas9 cells.

Mass spectrometry proteomics data have been deposited at the ProteomeXchange Consortium via the PRIDE partner repository ^86^ with the dataset identifier PXD060548.

---

**Supplementary Figure 1.**
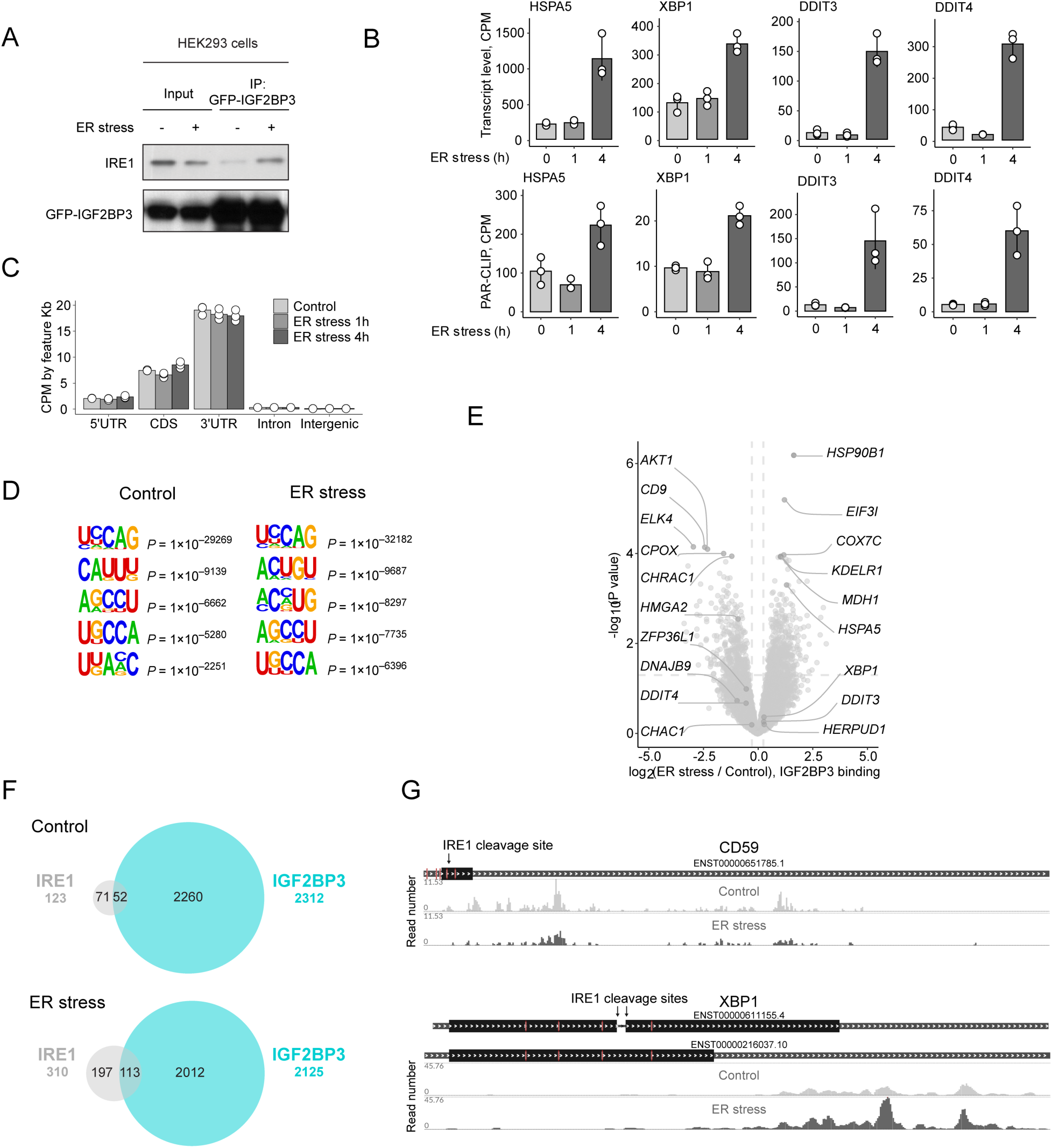
**A.** Western blot of IRE1 showing its association with IGF2BP3 after immunoprecipitation of GFP-IGF2BP3 from HEK293T cells treated with ER stress-inducing drug tunicamycin (TM) at 5 µg/mL for 4 hours. **B.** Barplots showing normalized read count numbers (CPM) for IGF2BP3-bound reads (IR-PAR-CLIP) and total transcript levels (QuantSeq) of selected UPR target genes upon ER stress induction with TM at 5 µg/mL for 1 and 4 hours. Values are the mean ± s.d of n=3 biological replicates. *P* values were calculated by two-sided Student’s t-test. **C.** Feature length-normalized aggregated IGF2BP3 IR-PAR-CLIP coverage of genomic features. **D.** 5nt-long consensus motifs enriched in IGF2BP3 IR-PAR-CLIP reads over scrambled control in control and ER stress conditions (4 hr). Motif enrichment analysis was performed using HOMER software. **E.** Volcano plot showing changes in IGF2BP3 binding (IR-PAR-CLIP CPM / QuantSeq CPM) upon ER stress and control conditions. n=3 biological replicates. *P* values were calculated by edgeR glmQLFTest. **F.** Venn diagram showing the intersection between IGF2BP3 and IRE1 PAR-CLIP targets ^30^ in control and ER stress conditions. **G.** IGF2BP3 IR-PAR-CLIP coverage of IRE1 targets *XBP1* and *CD59*.

**Supplementary Figure 2.**
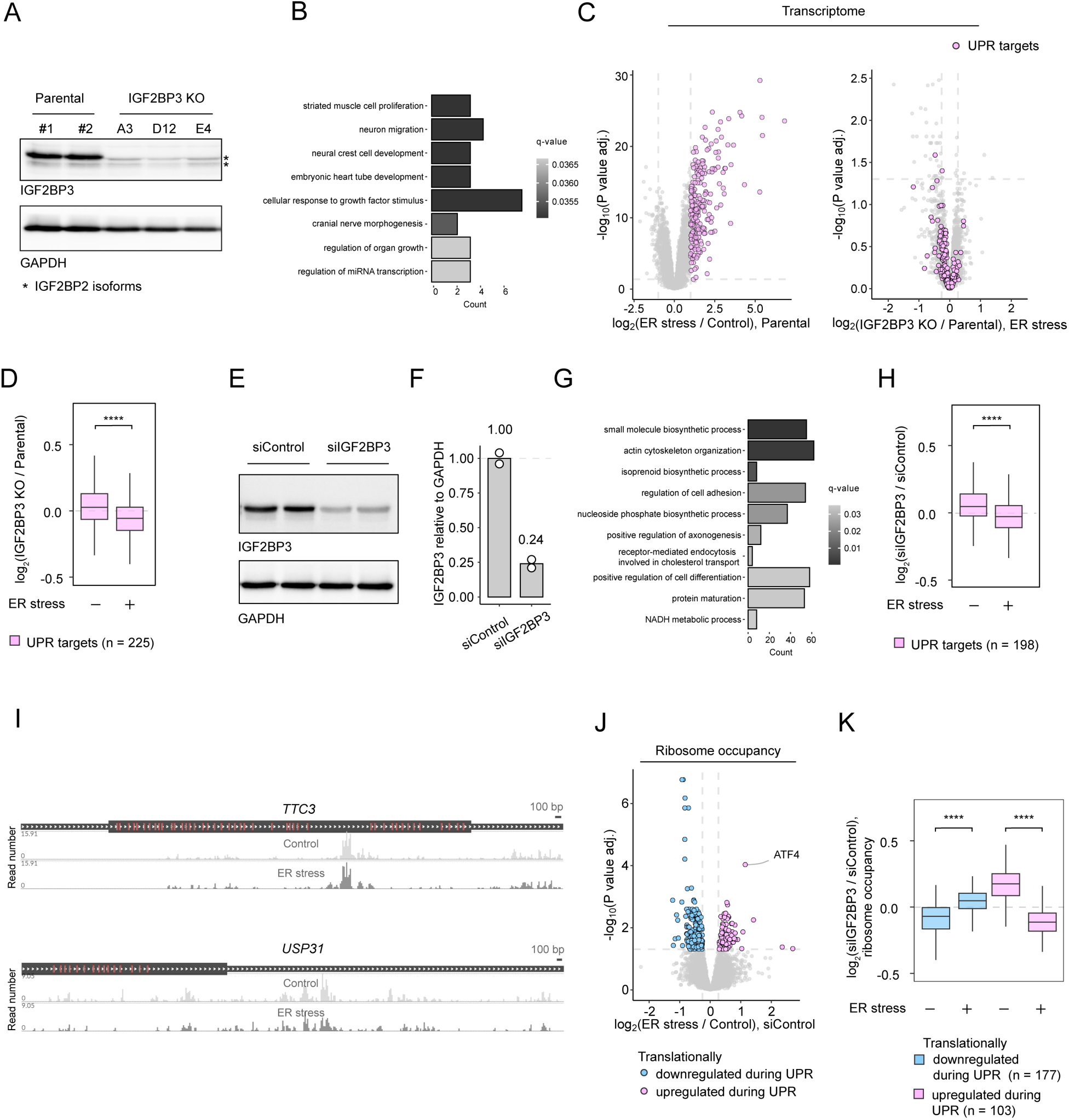
**A.** Western blot of parental and IGF2BP3 KO HCT116 cell lines (clones A3, D12 and E4). * marks IGF2BP2 isoforms recognized by polyclonal anti-IGF2BP3 antibody. **B.** GO term analysis of genes downregulated in IGF2BP3 KO comparing to parental cells during ER stress. edgeR glmQLFTest *P* value adj. < 0.05. **C.** Volcano plot of transcriptome (RNA-seq) changes upon ER stress treatment showing the selection of UPR targets (upregulated more two-fold upon ER stress)(left panel), and volcano plot upon IGF2BP3 KO in ER stress conditions with highlighted UPR targets (right panel). *P* values were calculated by edgeR glmQLFTest. **D.** Boxplot showing changes in transcript levels upon IGF2BP3 KO for UPR targets. *P* values were calculated by two-sided Wilcoxon test. **E.** Western blot showing depletion of IGF2BP3 after 48-hour treatment with siRNA against IGF2BP3 compared to control siRNA. **F.** Quantification of D. **G.** GO term analysis of genes downregulated upon siRNA-mediated depletion of IGF2BP3 comparing to siControl during ER stress. edgeR glmQLFTest *P* value adj. < 0.05. **H.** Boxplot showing changes in transcript levels upon siRNA-mediated depletion of IGF2BP3 for UPR targets. *P* values were calculated by two-sided Wilcoxon test. **I.** IGF2BP3 IR-PAR-CLIP coverage of *TTC3* and *USP31.* **J.** Volcano plot of ribosome occupancy (RO, log2(RiboSeq CPM / RNA-seq CPM)) changes upon ER stress showing the selection of translationally regulated UPR targets (ΔRO > 20% and *P* value adj. < 0.05). *P* values were calculated by edgeR glmQLFTest. **K.** Boxplot showing changes in ribosome occupancy upon siRNA-mediated IGF2BP3 depletion for UPR targets. *P* values were calculated by two-sided Wilcoxon test. \**P* < 0.05; \*\**P* < 0.01; \*\*\**P* < 0.001; \*\*\*\**P* < 0.0001.

**Supplementary Figure 3.**
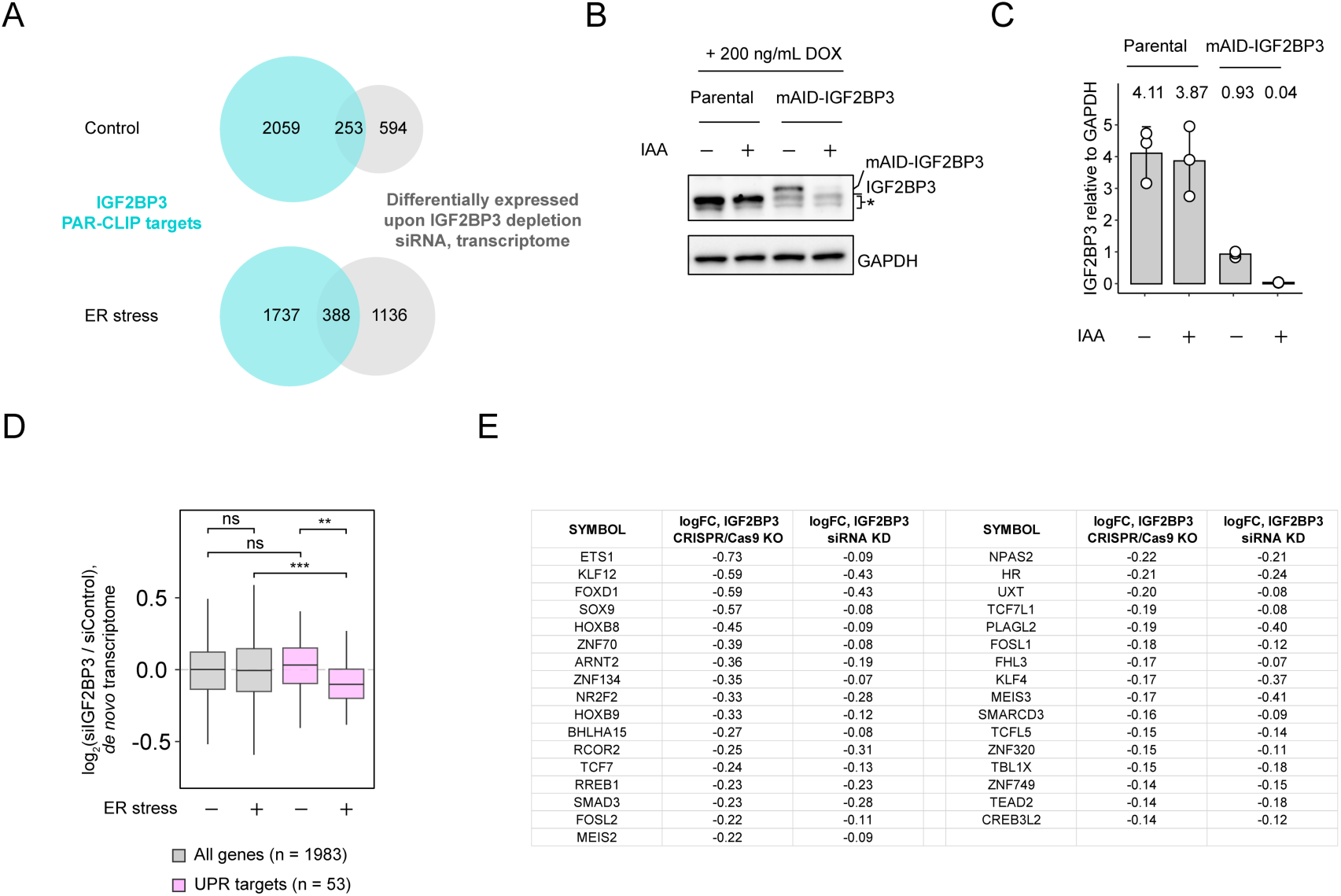
**A.** Venn diagram showing the intersection between IGF2BP3 targets and IGF2BP3-regulated genes (ΔRNA-seq > 20% and *P* value adj. < 0.05). **B**. Western blot showing auxin-induced depletion of mAID-IGF2BP3. * marks IGF2BP2 isoforms recognized by polyclonal anti-IGF2BP3 antibody. **C**. Quantification of **B** n=3 biological replicates. **D.** Boxplot showing changes in *de novo* transcripts levels of UPR targets (synthesized during 2-hour s^4^U pulse) T-C(SLAMseq T-C CPM) upon siRNA-mediated depletion of IGF2BP3. *P* values were calculated by two-sided Wilcoxon test. \**P* < 0.05; \*\**P* < 0.01; \*\*\**P* < 0.001; \*\*\*\**P* < 0.0001. **E.** Table showing log_2_(IGF2BP3 depletion / control) values for transcriptional regulators (GO:0140110) that were downregulated more than 10% or 5% upon IGF2BP3 CRISPR/Cas9- or siRNA-mediated depletion, respectively, under ER stress conditions.

**Supplementary Figure 4.**
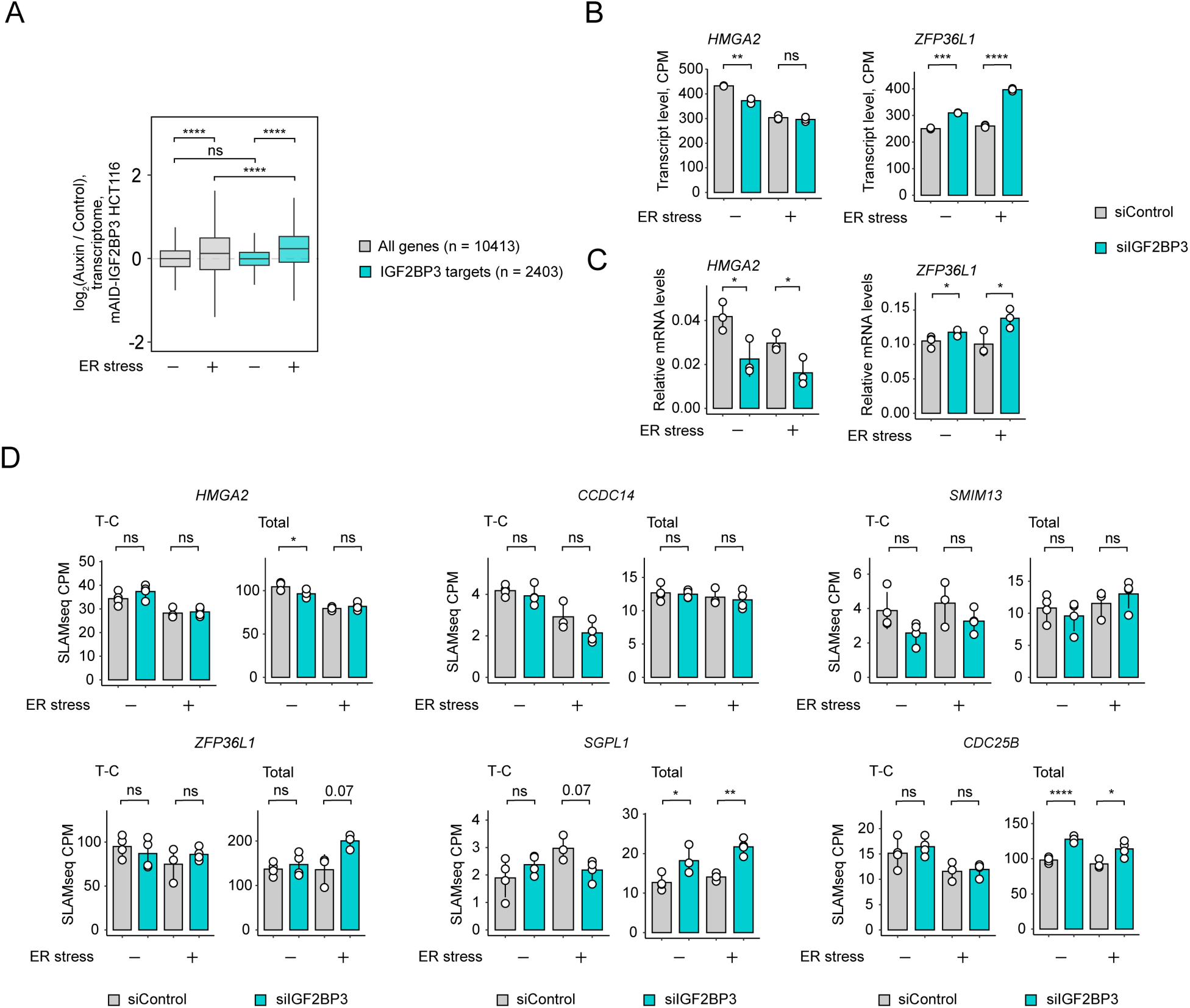
**A.** Boxplot showing changes in total transcript levels upon auxin-induced mAID-IGF2BP3 depletion for all genes and IGF2BP3 targets control and ER stress conditions. *P* values were calculated by two-sided Wilcoxon test. **B.** Barplots showing total RNA-seq CPM values normalized to siControl conditions for *HMGA2* and *ZFP36L1*. Values are the mean ± s.d of n=3 biological replicates. *P* values were calculated by paired two-sided Student’s t-test. **C.** Barplots showing results of RT-qPCR analyses of *HMGA2* and *ZFP36L1* (mRNA levels relative to *RPL6*) in unstressed cells and cells treated with tunicamycin at 5 µg/mL for 6 hours upon siRNA-mediated depletion of IGF2BP3. Values are the mean ± s.d. of n=3 biological replicates. *P* values were calculated by two-sided Student’s t-test. **D.** Barplots showing SLAMseq total (total CPM) and *de novo* (T-C CPM) transcript levels for selected transcripts regulated by IGF2BP3. Values are the mean ± s.d of n=4 biological replicates. *P* values were calculated by two-sided Student’s t-test. \**P* < 0.05; \*\**P* < 0.01; \*\*\**P* < 0.001; \*\*\*\**P* < 0.0001.

**Supplementary Figure 5.**
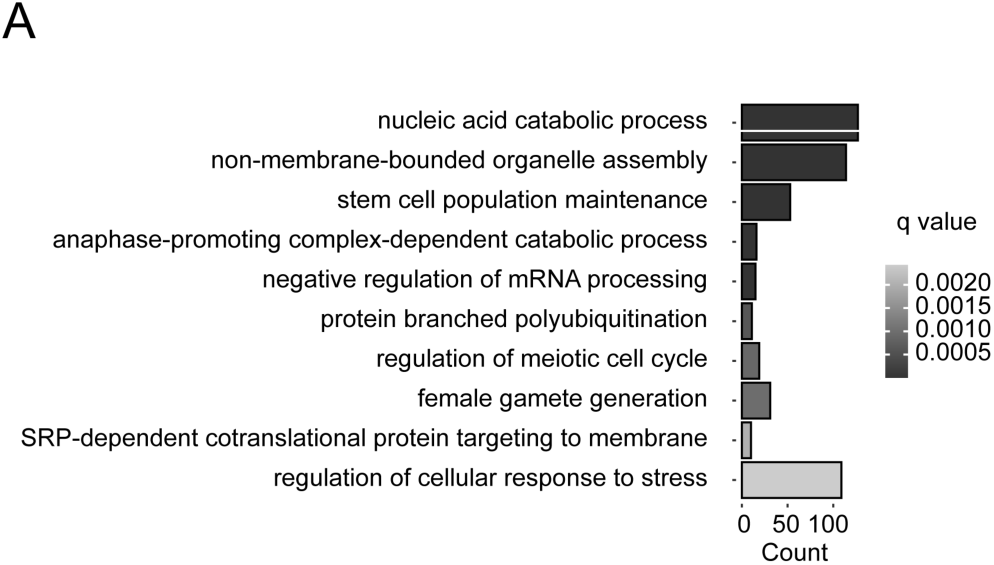
**A.** GO term enrichment analysis for IGF2BP3 interaction partners (4 times enriched over IgG control, *P* value adj. < 0.05).

## Notes

### Competing Interest Statement

S.L.A. declare competing interests based on a granted patent related to SLAMseq. S.L.A. is co-founder, advisor, and member of the board of QUANTRO Therapeutics GmbH.

